# Hydrogel-imposed boundary conditions guide single-lumen neuroepithelial morphogenesis

**DOI:** 10.64898/2026.01.30.702717

**Authors:** Michelle S. Huang, Julien G. Roth, Dohui Kim, Kristine P. Pashin, Dominic Pizzarella, Tianming M. Yang, Yueming Liu, Renato S. Navarro, Theo D. Palmer, Sarah C. Heilshorn

**Affiliations:** Department of Chemical Engineering, Stanford University, Stanford, CA, USA; The Institute for Chemistry, Engineering & Medicine for Human Health (Sarafan ChEM-H), Stanford University, Stanford, CA, USA; Institute for Stem Cell Biology and Regenerative Medicine, Stanford University School of Medicine, Stanford, CA, USA; Complex In Vitro Systems, Translational Safety, Genentech Inc., South San Francisco, CA, USA; Department of Materials Science and Engineering, Stanford University, Stanford, CA, USA; Department of Bioengineering, Stanford University, Stanford, CA, USA; Department of Materials Science and Engineering, University of Florida, Gainesville, FL, USA; Department of Neurosurgery, Stanford University School of Medicine, Stanford, CA, USA

## Abstract

Three-dimensional (3D) stem cell-based cultures have emerged as promising *in vitro* model systems for studying human neurodevelopment. Current neural organoid protocols lack well-defined extracellular matrix (ECM) signaling and are limited by the formation of irregular tissue morphologies with multiple organizing centers, in contrast to the single neuroepithelial structure that emerges during embryonic development. This variability limits inter-organoid reproducibility and constrains their utility for modeling early developmental processes. To overcome these limitations, we leverage a materials-based approach to impose dynamic boundary conditions that extrinsically guide the self-organization of human induced pluripotent stem cells (iPSCs). Specifically, we develop a family of hyaluronic acid–elastin-like protein (HELP) hydrogels crosslinked with dynamic covalent bonds that recapitulate key biochemical and biophysical properties of the developing human neural ECM. Within these HELP hydrogels, iPSCs robustly self-organize from a single cell into complex neuroepithelial tissues with a single lumen. By tuning the boundary conditions imposed by the hydrogel, we identify matrix stress relaxation rate and tensional homeostasis as key regulators of single-lumen rosette formation and maintenance. With this hydrogel-enabled system, we identify phenotypic abnormalities in an early neurodevelopmental model of 22q11.2 deletion syndrome. Ultimately, our tunable engineered hydrogel supports the initiation of single-cell derived 3D neuroepithelial tissues, enables investigation into how matrix-imposed boundary conditions guide developmental morphogenesis, and establishes a reproducible platform for disease modeling.

## Introduction

Neurodevelopment begins with the formation of a single neural tube, which involves the careful coordination of multiple cells simultaneously undergoing conserved morphogenetic programs. These include the establishment of neuroepithelial cell apical-basal polarity, proliferation at the apical surface of the nascent neural tube, and radial migration and differentiation at the basal surface.^1,2^ Dysregulated neural tube formation during development results in congenital malformations, such as diplomyelia, diastematomyelia, and anencephaly.^3^ Thus, establishing a singular neuroepithelium is a defining cytoarchitectural milestone for modeling early human brain development *in vitro*.

Recent advances in pluripotent stem cell (PSC)-derived neural organoids have enabled scalable models that recapitulate several features of the developing brain, including its cell type diversity, transcriptional profiles, and aspects of functional maturation.^4–8^ However, conventional organoid protocols commonly yield irregular tissue morphologies and multiple neuroepithelial units (or neural rosettes) rather than a single neuroepithelium, limiting their reproducibility and physiological accuracy.^9^ Approached from an engineering perspective, PSCs form a dynamic, self-organizing system that responds to its surrounding boundary conditions as inputs to guide morphogenesis.^10–14^ Organoid systems that lack appropriate constraints are therefore underconstrained; that is, without defined extrinsic guidance, multiple morphogenetic outcomes are equally permissive, yielding variable and often disorganized structures.

As one strategy to provide constraints to the emerging structure, micropatterning approaches have demonstrated that restricting initial cell placement and spatial orientation can direct neural tube-like tissue formation.^15–17^ Similarly, other top-down engineering strategies, which include microwell plates,^18–20^ geometrically patterned materials,^21–24^ or physical microdissection of self-organizing cell assemblies,^25–27^ have been used to impose defined geometric constraints that enable reproducible cell patterning and organoid formation. These methods all rely on experimenter-imposed boundary conditions, however, *in vivo* these constraints arise naturally from the extracellular matrix (ECM). We therefore hypothesized that the incorporation of an engineered hydrogel could impose the initial boundary conditions that would guide and stabilize early morphogenetic programs *in vitro*, resulting in reproducible, predictable structures that self-emerge without additional top-down constraints.

To achieve this goal, we take a bioinspired materials-based approach to introduce matrix-imposed boundary conditions that guide the early stages of human brain morphogenesis. To date, a limited number of studies have explored the use of biomaterials to intentionally drive neurodevelopmental programs in 3D cellular models. A few such studies have encapsulated pre-formed PSC aggregates in hydrogels, but these approaches introduced matrix cues after critical polarity-establishing events have already occurred.^28–30^ Alternatively, early dynamic mechanical stimulation in the form of top-down actuation supported neuroepithelial self-patterning and polarity acquisition.^31^ We reasoned that a biomimetic approach to exerting dynamic mechanical properties could be realized through hydrogel design. Specifically, we leverage a viscoelastic hydrogel with tunable stress relaxation kinetics to impose and probe the effects of dynamic boundary conditions. In previous work by us and others, hydrogels have been explored as a means to provide matrix-derived signaling cues that can guide neural differentiation at the length-scale of individual cells.^32–36^ Here, we propose that exerting engineering control at the earliest stages of self-organization is essential for the emergence of a single, well-organized neuroepithelium.

In this work, we develop a family of protein-engineered hydrogels consisting of hyaluronic acid and elastin-like protein crosslinked with dynamic covalent bonds that imparts neural ECM-derived biochemical and biophysical signaling cues to guide the self-organization of 3D neural rosettes. Within this fully defined synthetic hydrogel, human PSCs robustly form neuroepithelial tissues containing a single neural rosette emerging from a single PSC. We leverage the tunability of our engineered hydrogel to evaluate how altering boundary conditions destabilizes or restricts morphogenetic outcomes. Lastly, we demonstrate that 3D single-lumen neural rosettes can robustly manifest genetically-driven phenotypes of 22q11.2 deletion syndrome. Ultimately, by combining cell-intrinsic morphogenetic momentum with extrinsically imposed material boundary conditions, we establish a reproducible and scalable approach for generating single-lumen neuroepithelial tissues suitable for mechanistic studies of development and disease modeling.

### Hydrogel boundary conditions guide single-lumen neural rosette formation

To impose boundary conditions that can guide lumen formation in 3D, we began by designing an engineered hydrogel that emulates aspects of the *in vitro* brain microenvironment. We sought to incorporate biochemical signaling from ECM components that are present during neurodevelopment, including hyaluronic acid (HA) and fibronectin.^37^ In particular, HA has been implicated in neuroepithelial expansion and migration,^38^ and fibronectin, which is enriched in the surrounding non-neural ectoderm, plays a crucial role in neurulation.^16^ Therefore, we designed our engineered hydrogel to consist of recombinant HA and elastin-like protein (ELP) crosslinked to form a HELP hydrogel (**Supplementary Fig. 1a**).^39^ The ELP used here is a recombinantly-expressed engineered protein that contains alternating elastin-derived and fibronectin-derived motifs, which include the integrin-binding RGD cell-adhesive ligand (**Supplementary Fig. 1a**). The mechanical properties of the HELP hydrogel were tuned to match the stiffness of native brain tissue (∼800 Pa; **Supplementary Fig. 1b,c**).^33^ For the crosslinks, we selected hydrazone bonds, which are a type of dynamic covalent chemistry (DCC). The reversible nature of the dynamic covalent bond results in a stress-relaxing hydrogel that permits cell-mediated matrix remodeling (**Supplementary Fig. 1d**). In previous work, HELP hydrogels with this stress relaxation rate enabled the volumetric expansion of single-lumen human enteroids from single intestinal stem cells;^40^ thus, we hypothesized that this combination of defined biochemical cues and mechanical resistance provided by the matrix would support the emergence of 3D single-lumen neural rosettes.

First, we evaluated how the boundary conditions imposed by our hydrogel system influenced rosette formation compared to a condition lacking imposed boundaries, where iPSCs self-organize freely in suspension. In the suspension protocol, which mirrors the early stages of conventional guided neural organoid protocols,^5,41^ iPSCs are first aggregated and then maintained in a suspension culture devoid of exogenous ECM components (**Fig. 1a**). In the HELP protocol, iPSCs are embedded as single cells within the hydrogels and allowed to proliferate and self-organize (**Fig. 1a**). In both protocols, the same neural induction factors were introduced at the same time (day 2), including a TGF-β inhibitor (SB-431542) and a BMP pathway inhibitor (LDN-193189). By day 8 (after six days of neural induction) of the suspension protocol, iPSCs had already formed multiple rosettes in each spheroid (**Fig. 1b**), consistent with previous reports of the non-physiological emergence of multiple organizing centers within more mature neural organoids.^5,42,43^ In addition, the size and number of rosettes per spheroid that emerged in the suspension protocol exhibited high variability (**Fig. 1b,c**). In contrast, within engineered HELP hydrogels, single-cell iPSCs had self-organized into multicellular neuroepithelial units each containing a single lumen with significantly greater uniformity in size (**Fig. 1b,c**). Immunostaining revealed that these neural rosettes have acquired apical-basal polarity, with Sox2+ neuroepithelial cells radially arranged around a single lumen marked by N-cadherin and a characteristic actomyosin contractile ring at the apical end feet (**Fig. 1d**).

**Figure 1.**
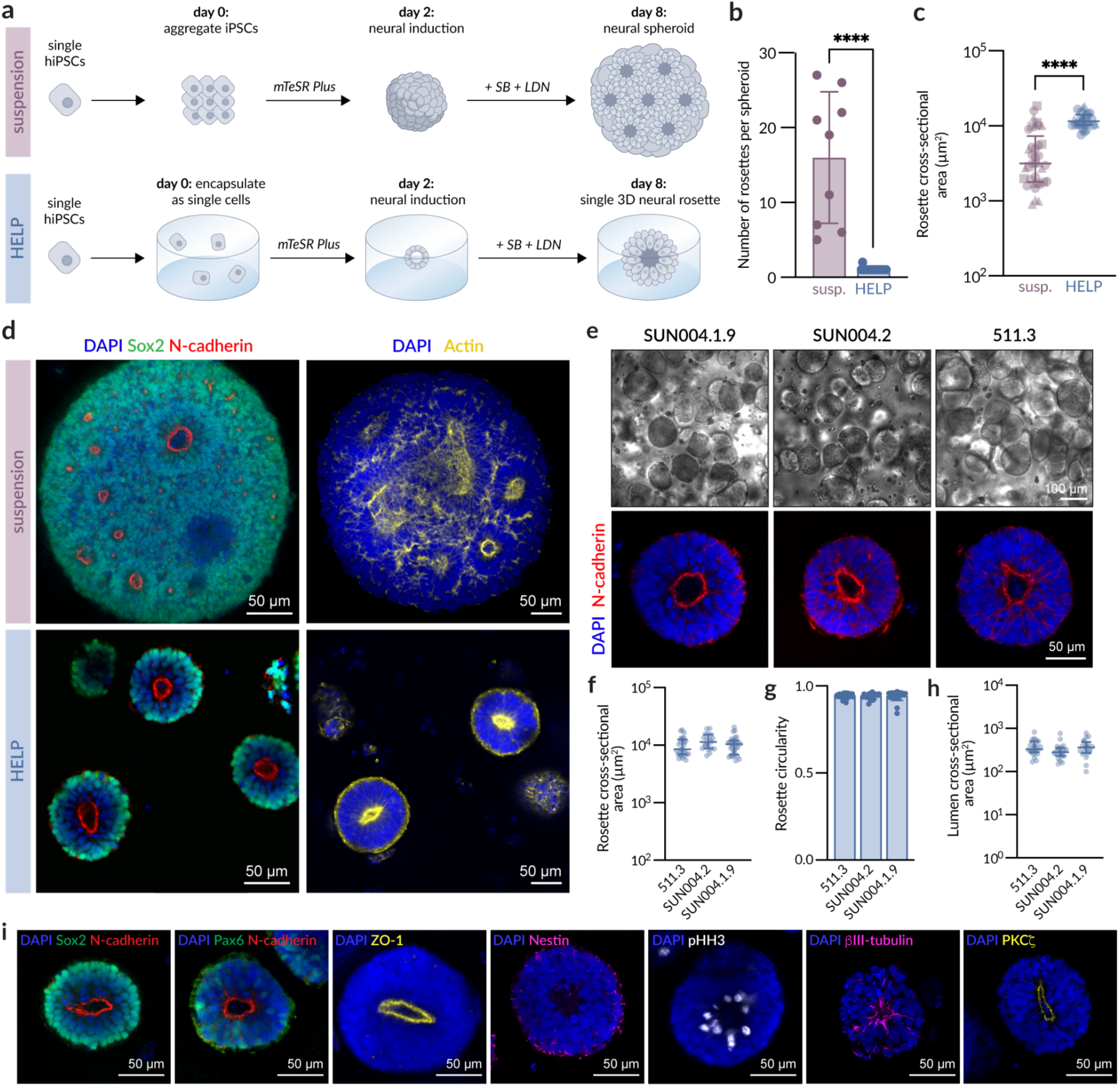
hiPSCs form single-lumen 3D neural rosettes within engineered HELP hydrogels. **a.** Schematic of experimental workflow to generate multi-lumen neural spheroids in suspension (top) and single-lumen neural rosettes in engineered hydrogels (bottom). **b.** Quantification of the number of rosettes per spheroid at day 8 (suspension, *n* = 9 spheroids; HELP, *n* = 18 spheroids; data are means ± standard deviation). **c.** Quantification of rosette cross-sectional area in multi-lumen spheroids compared to single-lumen neural rosettes at day 8 (suspension, *N* = 3 independent spheroids, *n* = 37 rosettes; HELP, *N* = 3 independent experimental replicate hydrogels, *n* = 28 rosettes; data are medians with interquartile range). **d.** Representative fluorescence images of multi-lumen neural spheroids in suspension and single-lumen neural rosettes in engineered hydrogels at day 8 labeled with Sox2 (green) and N-cadherin (red). Nuclei are counterstained with DAPI (blue), and F-actin is counterstained with phalloidin (yellow). **e.** Representative brightfield (top) and fluorescence (bottom) images of single-lumen neural rosettes in engineered hydrogels at day 8 for multiple iPSC lines labeled with N-cadherin (red) and a counterstain for nuclei (DAPI, blue). **f.** Quantification of rosette cross-sectional area for multiple iPSC lines at day 8 (*N* = 3 independent experimental replicate hydrogels; 511.3, *n* = 27 rosettes; SUN004.2, *n* = 19 rosettes; SUN004.1.9, *n* = 29 rosettes; data are medians with interquartile range). **g.** Quantification of rosette circularity for multiple iPSC lines at day 8 (*N* = 3 independent experimental replicate hydrogels; 511.3, *n* = 27 rosettes; SUN004.2, *n* = 19 rosettes; SUN004.1.9, *n* = 29 rosettes; data are means ± standard deviation). **h.** Quantification of lumen cross-sectional area for multiple iPSC lines at day 8 (*N* = 3 independent experimental replicate hydrogels; 511.3, *n* = 23 rosettes; SUN004.2, *n* = 21 rosettes; SUN004.1.9, *n* = 20 rosettes; data are medians with interquartile range). **i.** Representative fluorescence images of single-lumen neural rosettes in engineered hydrogels at day 8 immunostained for the designated proteins. Statistical analyses were unpaired t-test (b), Mann-Whitney test (c), Kruskal-Wallis with Dunn’s multiple comparisons test (f, h), and one-way ANOVA with Tukey’s multiple comparisons test (g). ****p < 0.0001.

To ensure reproducibility, the results were validated in six different iPSC lines from four donors, accounting for both intra- and inter-patient heterogeneity. Across all six lines, the rosettes exhibited similar growth rates and were relatively uniform in both size and morphology (**Fig. 1e-g; Supplementary Fig. 2a-c**). By day 8, polarized single-lumen neuroepithelia reproducibly emerged in all iPSC lines, with consistent lumen size across iPSC lines (**Fig. 1h**) and >90% of all spheroids containing a single lumen (**Supplementary Fig. 2a,d**).

Immunostaining was performed to validate neuroepithelial identity and the development of ventricular zone-like structures that emerge during early neural tube development. Day 8 single-lumen rosettes cultured in HELP contained cells expressing the neuroepithelial markers Sox2, Nestin, and paired box 6 (Pax6), which are indicative of a radial glial cell identity, surrounding a central apical lumen (**Fig. 1i**). The apical lumen is demarcated by N-cadherin adherens junctions, the tight junction protein zonula occludens-1 (ZO-1), and protein kinase C zeta (PKCζ), which together confirm the proper establishment of apical-basal polarity (**Fig. 1i**). Proliferative cells, marked by phospho-histone H3 (pHH3), were found primarily surrounding the apical surface of the rosette, similar to the localization of mitotic radial glia *in vivo* (**Fig. 1i**).^44^ Lastly, expression of neuron-specific βIII-tubulin suggests the emergence of early-born neurons by day 8 (**Fig. 1i**). Altogether, our data demonstrate that boundary conditions imposed through an engineered hydrogel enable the robust formation of single-lumen neuroepithelial tissues.

### Reproducible neuroepithelial morphogenesis requires defined boundary conditions

Given that hydrogel encapsulation regulated single-lumen neuroepithelium formation, we next investigated whether this morphogenetic outcome requires a matrix with defined boundary conditions. We compared our engineered HELP hydrogels to a commercially available Engelbreth-Holm-Swarm (EHS) matrix (*i.e.*, Matrigel, Cultrex, etc.), which has been routinely used in multiple established neural organoid protocols as an extrinsic matrix to support neuroepithelium formation and expansion.^26,30,45,46^ These matrices have also been used to induce a transition from 2D monolayers to 3D lumen-containing structures as *in vitro* neural tube models.^15,16^ EHS matrices are animal-derived, reconstituted basement membrane mixtures known to contain a myriad of matrix components that support organoid growth, but they are poorly defined, lack tunability, and suffer from a high degree of batch-to-batch variability.^47^ In contrast, our recombinant HELP hydrogels are xeno-free, exhibit a wide range of biochemical and biophysical tunability, and are highly reproducible.^32,48,49^ Therefore, we hypothesized that our defined HELP hydrogels would better support reproducible single-lumen neural rosette formation compared to the standard EHS matrices.

We directly compared 3D neural rosette formation in HELP hydrogels and Matrigel, a representative EHS matrix. As above, single iPSCs were encapsulated in each hydrogel formulation, and neural induction factors were introduced on day 2. In Matrigel, although the iPSC clusters start out as individual spherical units, over time they begin to develop irregular, highly heterogeneous morphologies (**Fig. 2a**). This observation is consistent with previous reports that show the addition of Matrigel induces neuroepithelial budding and tissue irregularities.^50^ By day 8, when single-lumen neural rosettes have emerged in HELP hydrogels, those in Matrigel showed markedly less organized epithelial structures (**Fig. 2a**). Of note, immunostaining revealed N-cadherin to be distributed selectively at the apical surface for neural rosettes in HELP, whereas it displayed a more diffuse expression profile throughout the neural rosettes that formed in Matrigel (**Fig. 2b**). Also in HELP, Sox2 is expressed throughout the rosette but with higher intensity at the basal surface, which is consistent with the interkinetic nuclear migration of cycling neuroepithelial progenitors (**Fig. 2b; Supplementary Fig. 3a**).^51,52^ In contrast, Sox2 expression in Matrigel is relatively diffuse throughout the tissue, with higher variability between rosettes, consistent with a reduction in radial glial polarity and loss of clear apical-basal tissue organization.

**Figure 2.**
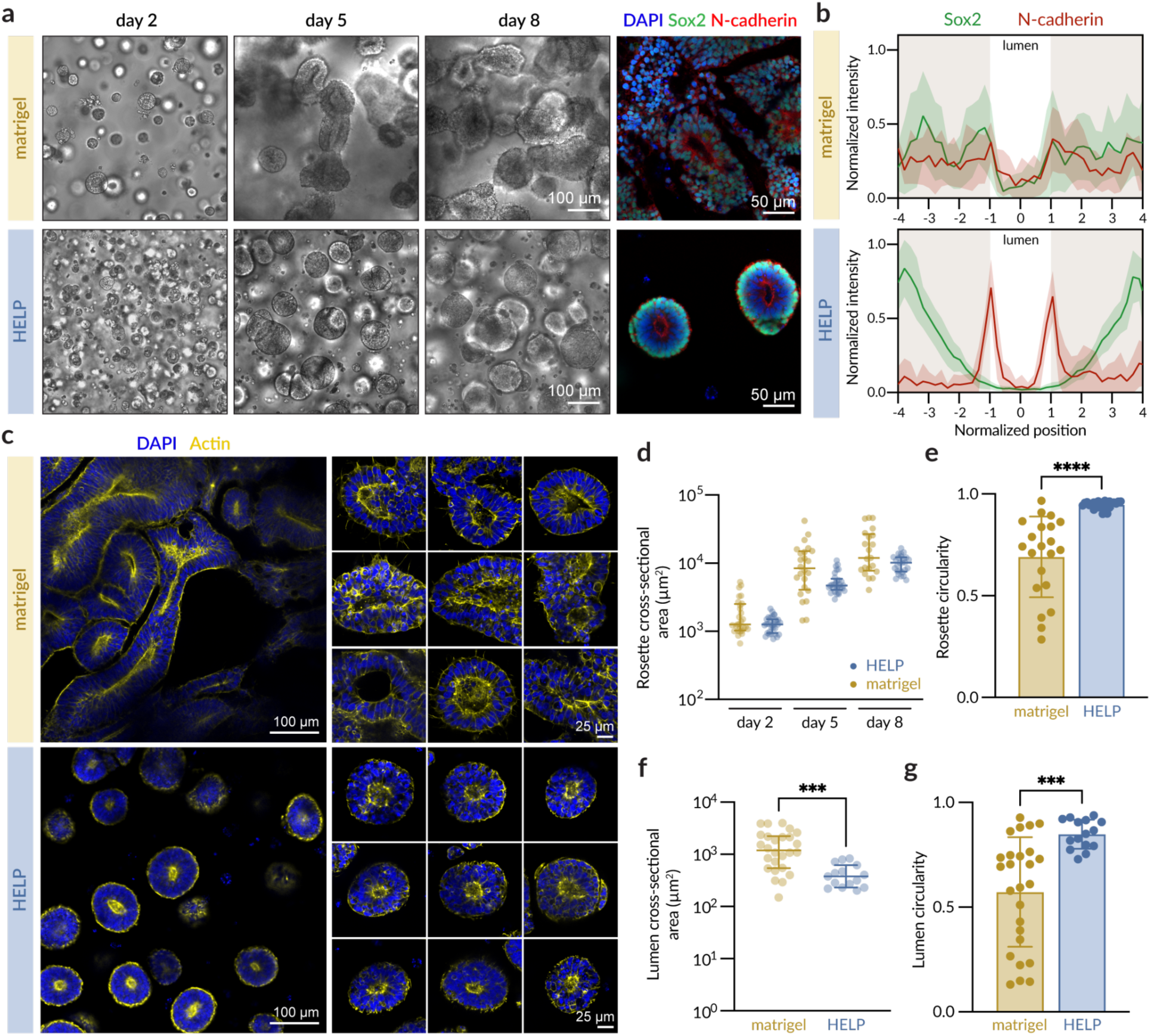
Defined hydrogel boundary conditions enable reproducible formation of 3D single-lumen neural rosettes. **a.** Representative brightfield images at days 2, 5, and 8 (left) of neural rosettes in Matrigel and HELP formed by encapsulating individual cells at day 0 and introducing neural induction factors at day 2. Fluorescence images of neural rosettes at day 8 (right) labeled with Sox2 (green) and N-cadherin (red). Nuclei are counterstained with DAPI (blue). **b.** Quantification of Sox2 and N-cadherin localization at day 8 in Matrigel and HELP. The position was normalized by placing the lumen center at 0 and the apical surfaces at −1 and +1 (Matrigel, *n* = 10 rosettes; HELP, *n* = 10 rosettes; the curve represents the mean and the shaded area represents the standard deviation). **c.** Representative fluorescence images at day 8 of neural rosettes in Matrigel and HELP showing nuclei (DAPI, blue) and F-actin (phalloidin, yellow). **d.** Quantification of rosette cross-sectional area at days 2, 5, and 8 (*N* = 3 independent experimental replicate hydrogels; Matrigel, *n* = 21-28 rosettes; HELP, *n* = 26-31 rosettes; data are medians with interquartile range). **e.** Quantification of rosette circularity at day 8 (*N* = 3 independent experimental replicate hydrogels; Matrigel, *n* = 21 rosettes; HELP, *n* = 26 rosettes; data are means ± standard deviation). **f.** Quantification of lumen cross-sectional area at day 8 (*N* = 3 independent experimental replicate hydrogels; Matrigel, *n* = 26 rosettes; HELP, *n* = 15 rosettes; data are medians with interquartile range). **g.** Quantification of lumen circularity at day 8 (*N* = 3 independent experimental replicate hydrogels; Matrigel, *n* = 26 rosettes; HELP, *n* = 15 rosettes; data are means ± standard deviation). Statistical analyses were Kruskal-Wallis with Dunn’s multiple comparisons test (d) and unpaired t-test (e, f, g). ***p < 0.001 and ****p < 0.0001.

Although some single-lumen neural rosettes developed in Matrigel, they were far less reproducible compared to those that formed within HELP hydrogels (**Fig. 2c**). Across several morphometrics, including rosette cross-sectional area (**Fig. 2d**), rosette circularity (**Fig. 2e**), lumen cross-sectional area (**Fig. 2f**), and lumen circularity (**Fig. 2g**), neural rosettes within HELP exhibited much less variance compared to those within Matrigel. Interestingly, even neural rosettes that grew adjacent to one another did not fuse within HELP hydrogels (**Supplementary Fig. 3b**). Thus, single-lumen neuroepithelia form reliably through self-assembly without the need for micropatterning or bioprinting to pre-specify cell positions. Importantly, single-lumen neural rosettes could be found distributed throughout the depth of the 3D hydrogel (**Supplementary Fig. 3c**), which here were 800 µm thick, offering greater scalability compared to 2D micropatterning approaches. Collectively, these data suggest that the specific hydrogel boundary conditions of our engineered HELP matrix support reproducible single-lumen neural rosette formation.

### Matrix viscoelasticity regulates 3D single-lumen neural rosette formation

To better understand what matrix properties govern single-lumen rosette formation, we next altered the boundary conditions imposed by our HELP hydrogels. In particular, we focused on tuning matrix viscoelasticity, as we and others have demonstrated that viscoelasticity has a profound effect on both neural cell fate acquisition and multicellular morphogenesis.^14,32,34,53,54^ When considering multicellular self-organization, we propose that the hydrogel matrix should not only be remodelable to facilitate multicellular volumetric expansion, but the matrix must also provide sufficient mechanical resistance to stabilize the emerging structure. That is, the material should exert dynamic confinement that supports the transfer of mechanical pressure from the material to the proliferating, self-organizing cells, yet still allows for remodeling of the matrix as the tissues grow.

We leveraged the tunability of our engineered HELP platform to alter matrix viscoelasticity independently of other boundary conditions (such as matrix stiffness, polymer concentration, and cell-adhesive ligand composition). Specifically, we engineered a family of HELP hydrogels with a range of stress relaxation rates by changing the identity of the covalent crosslinks (**Fig. 3a**). The previous experiments had been performed with a benzaldehyde-functionalized HA and a hydrazine-functionalized ELP to yield a viscoelastic, slow stress-relaxing HELP hydrogel (now referred to as Dyn. Slow HELP). By incorporating an aldehyde-functionalized HA, which has faster on/off rates, we designed a viscoelastic, fast stress-relaxing HELP hydrogel (referred to as Dyn. Fast HELP). By replacing the dynamic covalent crosslinks with static covalent crosslinks (tetrazine-functionalized HA and norbornene-functionalized ELP), we designed a slower stress-relaxing HELP hydrogel (referred to as Static HELP). All HELP hydrogels had a similar stiffness of ∼800 Pa (**Fig. 3b; Supplementary Fig. 4**), but they exhibited distinct stress relaxation profiles (**Fig. 3c**). The percentage of stress relaxed after 12 hours was ∼8%, ∼26%, and ∼67%, for Static, Dyn. Slow, and Dyn. Fast, respectively (**Fig. 3d**). While all three hydrogels have stress relaxation rates that are slower than that reported for native brain tissue,^55^ this range of viscoelasticity is known to impact the proliferation rate and differentiation of neural progenitor cells.^32,33^

**Figure 3.**
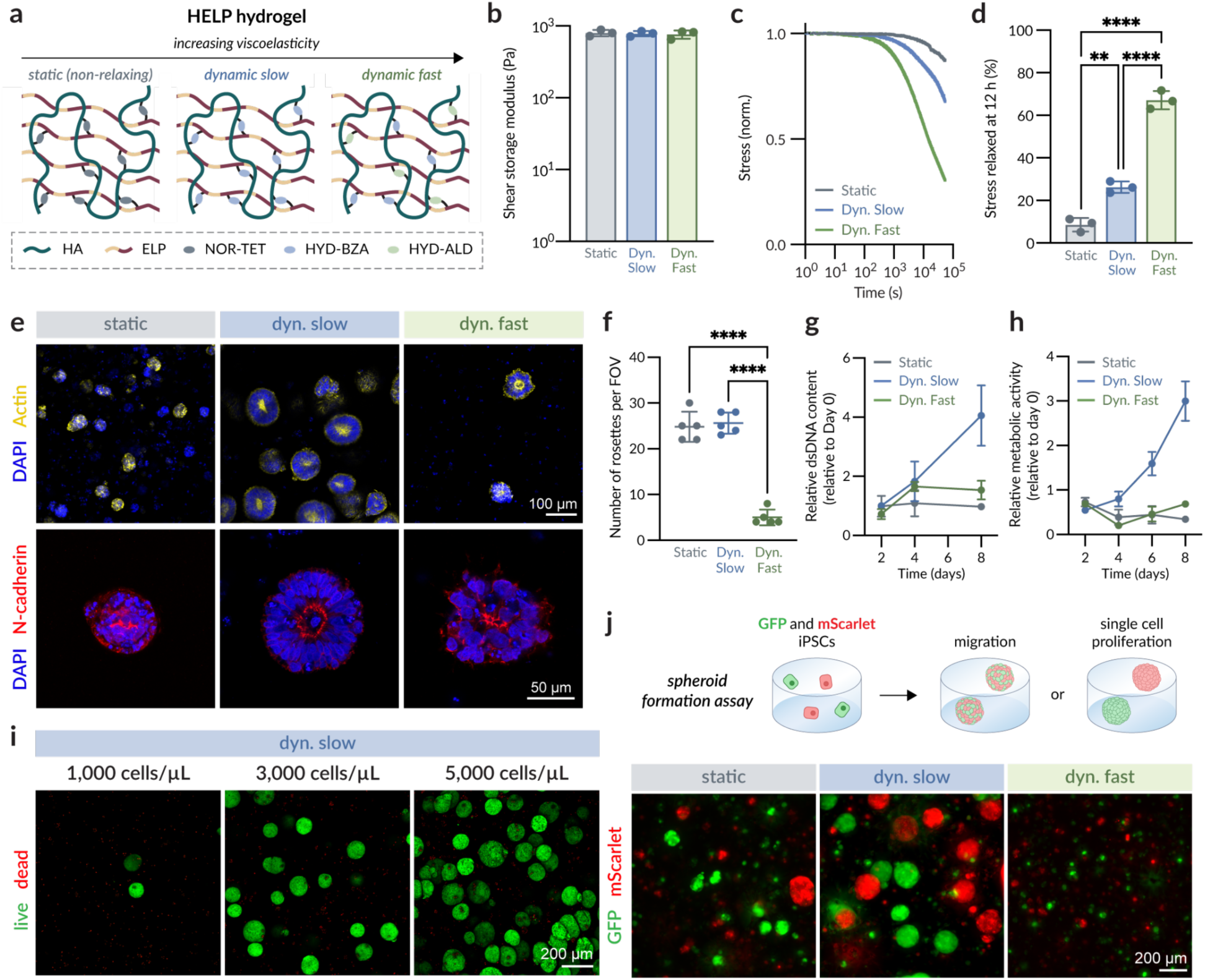
Matrix viscoelasticity regulates 3D neural rosette formation from encapsulated single cells. **a.** Schematic of Static, Dyn. Slow, and Dyn. Fast HELP hydrogels. **b.** Shear storage moduli of Static, Dyn. Slow, and Dyn. Fast HELP, reported at 1 rad/s (*N* = 3 independent experimental replicate hydrogels; data are means ± standard deviation). **c.** Representative stress relaxation curves of Static, Dyn. Slow, and Dyn. Fast HELP under a constant 10% strain. **d.** Percentage of initial stress relaxed after 12 h under a constant 10% strain (*N* = 3 independent experimental replicate hydrogels; data are means ± standard deviation). **e.** Representative fluorescence images at day 8 of neural rosettes in Static, Dyn. Slow, and Dyn. Fast HELP labeled with N-cadherin (red) and counterstains for nuclei (DAPI, blue) and F-actin (phalloidin, yellow). **f.** Quantification of number of rosettes per field of view (FOV; *N* = 3 independent experimental replicate hydrogels; data are means ± standard deviation). **g.** Quantification of dsDNA content within bulk hydrogels (*N* = 4 independent experimental replicate hydrogels; data are means ± standard deviation). **h.** Quantification of metabolic activity from bulk hydrogels (*N* = 4 independent experimental replicate hydrogels; data are means ± standard deviation). **i.** Representative fluorescence images at day 8 of neural rosettes in Dyn. Slow HELP with initial cell seeding densities of 1,000 cells/µL, 3,000 cells/µL, and 5,000 cells/µL labeled with calcein-AM (live) and ethidium homodimer-1 (dead). **j.** Representative fluorescence images at day 8 of neural rosettes formed from the co-encapsulation of GFP- and mScarlet-expressing iPSCs in Static, Dyn. Slow, and Dyn. Fast HELP. Statistical analyses were one-way ANOVA with Tukey’s multiple comparisons test (b, d, f). **p < 0.01 and ****p < 0.0001.

With this family of designer hydrogels, we evaluated the effect of matrix viscoelasticity on single-lumen neural rosette formation. Encapsulated iPSCs exhibited distinct morphological differences within the different hydrogel formulations (**Fig. 3e**). In particular, multicellular iPSC clusters formed within Static HELP gels, but they were smaller and lacked proper internal organization compared to those grown in Dyn. Slow HELP gels. On the other hand, iPSC clusters in Dyn. Fast HELP gels acquired proper polarization but exhibited highly irregular morphologies. Of significant note, far fewer neural rosettes were formed in the Dyn. Fast HELP gels compared to both Static and Dyn. Slow conditions (**Fig. 3f**). Overall, the Dyn. Slow HELP gels best supported the formation and subsequent expansion of 3D neural rosettes, while maintaining proper apical-basal polarity (**Fig. 3e,g,h**).

To begin exploring the mechanisms that underlie these morphological differences, we asked whether single-lumen rosette formation within our HELP gels arises through cell migration or single-cell proliferation to form early cell clusters. In a 3D context, matrix viscoelasticity is known to regulate both cell migration and proliferation, with faster-relaxing matrices typically promoting greater cell motility and slower-relaxing matrices favoring spheroid proliferative expansion.^33,54,56,57^ We therefore hypothesized that the successful establishment of single-lumen rosettes within Dyn. Slow HELP gels is primarily due to cell proliferation rather than migration. To test this hypothesis, we first evaluated whether different cell seeding densities would alter rosette formation within Dyn. Slow HELP gels. As expected, a higher seeding density yielded more rosettes, but the individual rosettes remained similar in size, morphology, and organization (**Fig. 3i; Supplementary Fig. 5**). These results indicate that single-lumen rosette morphogenesis occurs independently of seeding density, which suggests that each neural rosette may be emerging from a single proliferating iPSC embedded within a HELP hydrogel.

To confirm that neural rosettes emerged from a single iPSC rather than a cluster of nearby iPSCs, we performed a spheroid formation assay in which GFP- and mScarlet-expressing iPSCs were co-encapsulated as single cells within the same HELP hydrogel. When encapsulated separately, both iPSC lines were able to form single-lumen neural rosettes with similar growth rates (**Supplementary Fig. 2**). When they were co-encapsulated, neural rosettes formed that were either entirely GFP-expressing or entirely mScarlet-expressing, rather than a mosaic spheroid of GFP- and mScarlet-expressing cells (**Fig. 3j**). This result confirmed that neural rosettes within HELP hydrogels emerge from a single iPSC, suggesting that boundary conditions exerted at the length-scale of a single cell guide multicellular morphogenesis.

### Matrix confinement impairs neuroepithelial expansion and self-organization

After identifying the hydrogel stress relaxation rate that optimally supports single-lumen neural rosette morphogenesis, we next evaluated what occurs at the cell–matrix interface underlying the observed morphological differences. Specifically, we explored the role of these hydrogel-imposed boundaries on early spheroid formation, a process that involves dynamic shape and volume change. In a 3D environment, these dynamic processes can be restricted by physical constraints imposed by the hydrogel boundaries. A hydrogel that stores elastic energy for longer times will have more resistance to network rearrangement, providing mechanical confinement to the 3D culture.^58–60^ Therefore, we hypothesized that a matrix that exerts too much confinement would hinder volumetric expansion, and in turn preclude proper self-organization and fate acquisition.

To evaluate if the Static HELP hydrogels were indeed confining cell growth, we performed time-lapse microscopy of encapsulated iPSCs. Live cell imaging began 24 hours after encapsulation, immediately after the ROCK inhibitor Y27632 was removed from the culture to avoid confounding effects. iPSCs in Static HELP gels remained spherical and exhibited limited change in size or morphology over 18 hours (**Fig. 4a; Supplementary Video 1**). In contrast, iPSC clusters in Dyn. Slow and Dyn. Fast HELP gels underwent dynamic morphological changes and isotropic expansion enabled by the viscoelastic remodelability of these matrices (**Fig. 4a; Supplementary Videos 2 and 3**). In addition, within Dyn. Fast HELP gels, some iPSC clusters were observed sending and retracting protrusions into and out of their surrounding matrix (**Supplementary Video 3; Supplementary Fig. 6a**), while none of the observed cultures in Dyn. Slow gels displayed this behavior.

**Figure 4.**
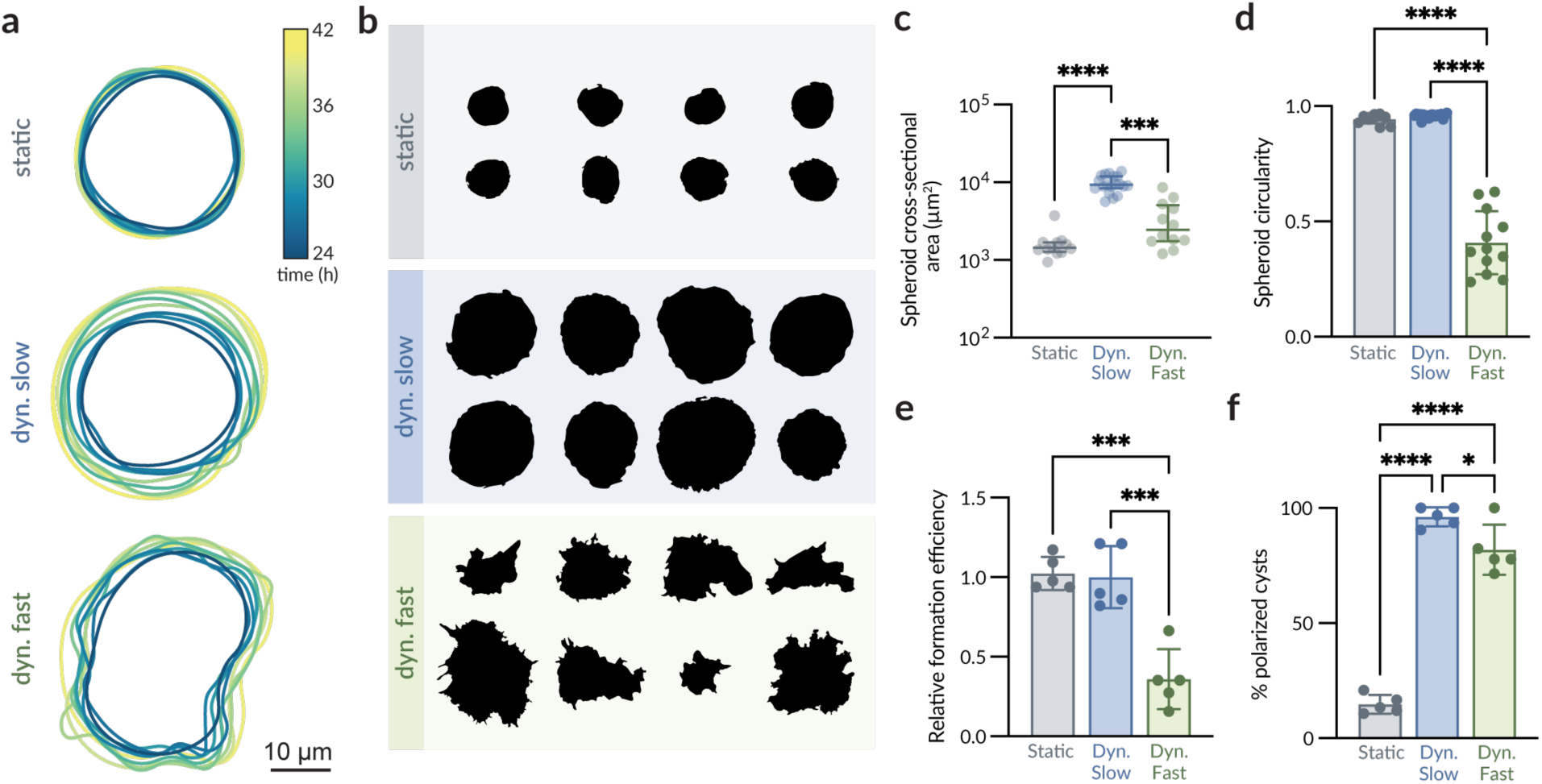
Matrix confinement impairs early volumetric expansion and self-organization. **a.** Overlaid outlines of the perimeter of iPSC clusters within Static, Dyn. Slow, and Dyn. Fast HELP from 24 h to 42 h of encapsulation. **b.** Representative masks of spheroids at day 8 within Static, Dyn. Slow, and Dyn. Fast HELP. **c.** Quantification of spheroid cross-sectional area at day 8 (*N* = 3 independent experimental replicate hydrogels; Static, *n* = 12 spheroids; Dyn. Slow, *n* = 19 spheroids; Dyn. Fast, *n* = 12 spheroids; data are medians with interquartile range). **d.** Quantification of spheroid circularity at day 8 (*N* = 3 independent experimental replicate hydrogels; Static, *n* = 12 spheroids; Dyn. Slow, *n* = 19 spheroids; Dyn. Fast, *n* = 12 spheroids; data are means ± standard deviation). **e.** Quantification of spheroid formation efficiency at day 8, normalized to Dyn. Slow (*N* = 5 independent experimental replicate hydrogels; data are means ± standard deviation). **f.** Percentage of apical-basally polarized cysts at day 8 based on actin staining (*N* = 5 independent experimental replicate hydrogels; data are means ± standard deviation). Statistical analyses were Kruskal-Wallis with Dunn’s multiple comparisons test (c) and one-way ANOVA with Tukey’s multiple comparisons test (d, e, f). *p < 0.05, ***p < 0.001, and ****p < 0.0001.

At day 8, the boundary conditions provided by the Static HELP matrix resulted in spheroids that were significantly smaller in size compared to those in Dyn. Slow and Dyn. Fast HELP matrices (**Fig. 4b,c**). The ability to form small iPSC clusters in Static HELP gels despite the matrix confinement is likely attributed to the relaxation of polymer entanglements, as the clusters eventually plateau in size. Interestingly, both the Static and Dyn. Slow HELP conditions yielded spheroids with high circularity (**Fig. 4b,d**), presumably due to the isotropic confinement that both of these hydrogels can provide. In contrast, in the less confining Dyn. Fast HELP gel, the resulting cellular structures at day 8 were highly irregular, as individual cells at the periphery of the spheroid extended into the hydrogel (**Fig. 4b,d**), consistent with the observations of dynamic cell protrusion on day 1. By day 8, while some cell clusters in Dyn. Fast exhibited volumetric expansion (**Fig. 4a**), others appeared to contract (**Supplementary Fig. 6a**). Overall, multicellular structures were much less likely to form at day 8 in the Dyn. Fast HELP gel compared to both Static and Dyn. Slow conditions (**Fig. 4e**).

We next examined how the ability to undergo early volumetric expansion correlated with internal cytoarchitecture and the acquisition of neuroepithelial cell identities at day 8. Notably, the smaller spheroids in the Static HELP gels yielded fewer apical-basally polarized structures compared to those in Dyn. Slow or Dyn. Fast gels (**Fig. 4e,f; Supplementary Fig. 6b**). The cells within these disorganized spheroids exhibited decreased expression of the neuroepithelial marker Pax6 at both the gene and protein levels (**Supplementary Fig. 6b,c**). Interestingly, even though the irregularly-shaped cell clusters in the Dyn. Fast gels were much less likely to form at day 8 (**Fig. 4e**), those that did form were about 4-times more likely to become polarized compared to cell clusters in the Static gels (**Fig. 4f**). Collectively, these results suggest that proper polarity and cell fate acquisition is preceded by early volumetric expansion, which is supported by a stress-relaxing hydrogel with remodelable crosslinks.

### Homeostatic tension is required for single-lumen rosette formation and maintenance

After establishing that early volumetric expansion on day 1 correlates with formation and polarization of a single lumen at day 8 in stress-relaxing hydrogels, we next asked which mechanisms may be involved by exploring an intermediate day 3 timepoint. Since the Static HELP gels were unable to reproducibly support volumetric expansion, that condition was not included in these subsequent studies. While initial iPSC viability 24 h post-encapsulation was high in both Dynamic HELP conditions, at day 3 there was a sharp decrease in viability for cells in Dyn. Fast HELP gels (**Fig. 5a**). This change in viability over time suggests that cell death was not caused by the encapsulation protocol, but rather occurred after matrix engagement.

**Figure 5.**
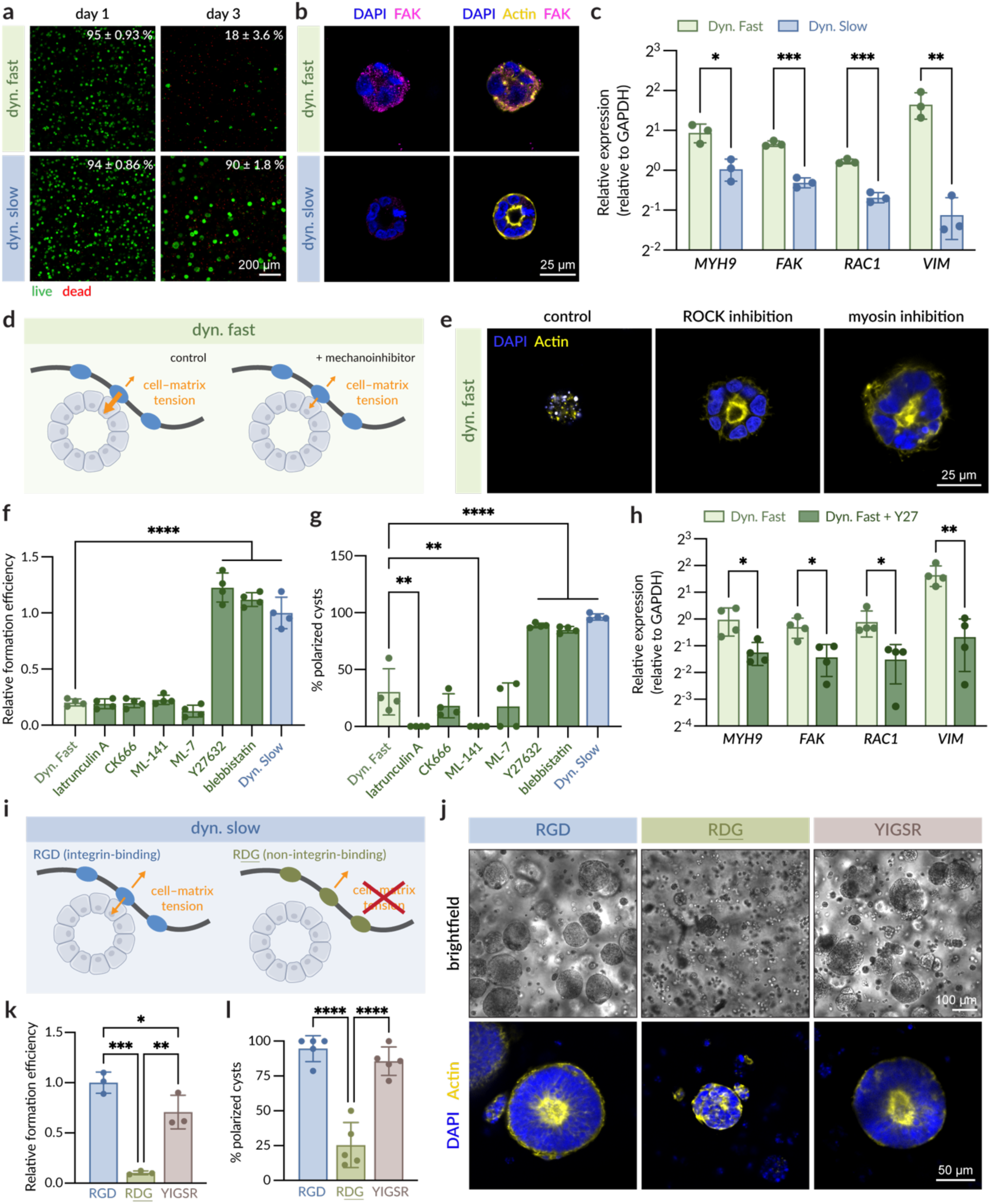
Tensional homeostasis enables single-lumen rosette formation in viscoelastic matrices. **a.** Representative fluorescence images at days 1 and 3 of neural rosettes in Dyn. Fast and Dyn. Slow HELP labeled with calcein-AM (live) and ethidium homodimer-1 (dead). Viability means and standard deviations are displayed (*N* = 3 independent experimental replicate hydrogels). **b.** Representative fluorescence images at day 3 of neural rosettes in Dyn. Fast and Dyn. Slow HELP labeled with focal adhesion kinase (FAK, magenta) and counterstains for nuclei (DAPI, blue) and F-actin (phalloidin, yellow). **c.** mRNA expression of *MYH9*, *FAK*, *RAC1*, and *VIM* after 48 h of culture (*N* = 3 independent experimental replicate hydrogels; data are means ± standard deviation). **d.** Schematic of proposed cell–matrix tension imbalance in Dyn. Fast HELP that is restored with mechanical tension inhibitors. **e.** Representative fluorescence images at day 3 of neural rosettes in Dyn. Fast HELP with 10 µM Y27632 or 10 µM blebbistatin showing nuclei (DAPI, blue) and F-actin (phalloidin, yellow). **f.** Quantification of spheroid formation efficiency with different mechanical inhibitors, normalized to Dyn. Slow (*N* = 4 independent experimental replicate hydrogels; data are means ± standard deviation). **g.** Percentage of apical-basally polarized cysts with different mechanical inhibitors (*N* = 4 independent experimental replicate hydrogels; data are means ± standard deviation). **h.** mRNA expression of *MYH9*, *FAK*, *RAC1*, and *VIM* after 48 h of culture in Dyn. Fast HELP with and without 10 µM Y27632 (*N* = 4 independent experimental replicate hydrogels; data are means ± standard deviation). **i.** Schematic of proposed hypothesis that removal of the cell-adhesive, integrin-binding RGD motif in Dyn. Slow HELP would upset tensional homeostasis. **j.** Representative brightfield images (top) and fluorescence images (bottom) of neural rosettes in Dyn. Slow HELP with different ligands showing nuclei (DAPI, blue) and F-actin (phalloidin, yellow). **k.** Spheroid formation efficiency in Dyn. Slow HELP with different ligands, normalized to RGD (*N* = 3 independent experimental replicate hydrogels; data are means ± standard deviation). **l.** Percentage of apical-basally polarized cysts in Dyn. Slow HELP (*N* = 5 independent experimental replicate hydrogels; data are means ± standard deviation). Statistical analyses were unpaired t-test (c, h), one-way ANOVA with Dunnett’s multiple comparisons test (f, g), and one-way ANOVA with Tukey’s multiple comparisons test (k, l). *p < 0.05, **p < 0.01, ***p < 0.001, and ****p < 0.0001.

To explore what matrix-induced signaling pathways may be dysregulated in Dyn. Fast HELP gels that lead to cell death, we next evaluated mechanobiological mechanisms that are known to be influenced by stress relaxation rate. We observed increased expression of several cytoskeletal tension-related markers in Dyn. Fast matrices compared to Dyn. Slow matrices (**Fig. 5b,c**). These include *MYH9*, the gene that encodes non-muscle myosin;^61^ *FAK*, the gene for focal adhesion kinase, which is involved in active integrin signaling;^62^ *RAC1*, the gene for a GTPase that is involved in mechanical signal transduction;^63^ and *VIM*, the gene for vimentin, an intermediate filament that regulates cell mechanics through actomyosin force transmission.^64^ Interestingly, at day 3, iPSC clusters in Dyn. Slow HELP gels were already polarized, while those in Dyn. Fast HELP gels exhibited elevated FAK expression and did not yet display apical-basal polarity (**Fig. 5b**). Together, these results led us to hypothesize that an imbalance in matrix tension within the Dyn. Fast HELP gels could lead to dysregulated mechanotransduction. In matrices that dissipate stress too rapidly, cells may be unable to maintain stable cell–matrix tension, resulting in signaling imbalance, loss of polarity cues, and eventual cell death.

To evaluate the hypothesis that a tension imbalance precludes polarity acquisition and rosette survival, we manipulated intracellular contractility within Dyn. Fast HELP gels by adding various small molecule inhibitors. If cellular tension exceeds the mechanical resistance provided by the matrix, this imbalance may disrupt tensional homeostasis; therefore, reducing actomyosin contractility should help restore equilibrium between cell-generated forces and matrix resistance (**Fig. 5d**). When actomyosin contractility was perturbed in Dyn. Fast HELP gels with Y27632 (ROCK inhibitor) or blebbistatin (myosin inhibitor), cell viability on day 3 was restored along with the formation of iPSC clusters, with similar size and single-lumen organization to the Dyn. Slow condition (**Fig. 5e,f; Supplemental Fig. 7a**). Notably, the iPSC clusters that formed in the presence of Y27632 and blebbistatin also acquired polarity similar to those in Dyn. Slow HELP gels (**Fig. 5e,g**). Inhibition of actin polymerization with latrunculin A, the Arp2/3 complex with CK666, Cdc42 GTPase with ML-141, and myosin light chain kinase with ML-7 had no effect on the ability to form iPSC clusters (**Fig. 5f**), with some inhibitors disrupting all actin organization entirely (**Fig. 5g; Supplementary Fig. 7b**). Additionally, the contractility-related genes that were previously upregulated in Dyn. Fast HELP gels showed reduced expression when Y27632 was added (**Fig. 5h**), suggesting that cellular mechano-dysregulation was reduced. Together, these results support the notion that restoring tensional homeostasis in a remodelable matrix is sufficient to enable rosette formation.

To test whether homeostatic tension with the matrix is required, we next disrupted cell–matrix adhesions in Dyn. Slow HELP gels by varying the cell-adhesive ligands within the protein-engineered ELP. By altering the amino acid sequence, we can replace the cell-adhesive RGD peptide with a scrambled, non-integrin-binding R<u>DG</u> peptide, resulting in mechanically-identical hydrogels with 0 mM RGD.^33^ By removing the ability for cells to pull on the matrix through RGD ligands, we hypothesized that this would disrupt cell–matrix tensional homeostasis (**Fig. 5i**). As expected, when iPSCs were encapsulated within R<u>DG</u>-presenting matrices, they exhibited significantly decreased formation efficiency and polarity acquisition (**Fig. 5j-l**). Finally, to evaluate whether single-lumen rosette formation is predicated on integrin–RGD interactions, we engineered an ELP displaying the laminin-derived, integrin-binding YIGSR motif. When iPSCs were encapsulated within YIGSR-presenting matrices, iPSCs were able to form rosettes despite the absence of RGD (**Fig. 5j-l**). Thus, in our engineered matrix system, the RGD and YIGSR ligands appear to be interchangeable, with either ligand capable of supporting early neuroepithelial morphogenesis. Altogether, these results suggest that single-lumen rosette formation is driven by integrin–matrix interactions, and that homeostatic tension is required for rosette formation.

### 22q11DS single-lumen neural rosettes from human iPSCs exhibit morphological abnormalities

22q11.2 deletion syndrome (22q11DS) is the most common chromosomal microdeletion in humans, affecting approximately 1 out of 3,000 live births.^65^ Individuals with this genetic deletion exhibit a range of neurodevelopmental phenotypes, including microcephaly, and an elevated risk for neuropsychiatric disorders (including schizophrenia and autism spectrum disorder) and intellectual disability. Rodent models of 22q11DS display abnormal neurogenesis and neuroanatomical differences including smaller brain sizes,^66–69^ suggesting an important role for the 22q11.2 locus in early brain development. While these rodent models have shed light on potential disease pathobiology, much of the human-specific developmental phenotypes remain underexplored. We reasoned that our hydrogel platform for reproducible generation of single-lumen neural rosettes from iPSCs would provide a tractable *in vitro* model to explore possible early neurodevelopmental differences with human, patient-specific 22q11DS samples.

To study the human-specific 22q11DS cultures in an early developmental context, we generated 3D single-lumen neural rosettes using iPSC lines from two individuals carrying the canonical 3-Mb 22q11.2 deletion.^70^ As with the healthy controls, both 22q11DS iPSC lines reproducibly formed multicellular units containing apico-basally polarized rosettes with similar formation efficiencies (**Fig. 6a-c; Supplementary Fig. 8a**). While the circular morphologies were maintained, the 22q11DS rosettes from both patient lines were notably smaller in size compared to control rosettes (**Fig. 6d; Supplementary Fig. 8b**). Importantly, this difference was only observed in HELP hydrogels and not in Matrigel, as both control and 22q11DS iPSCs in Matrigel developed irregular, heterogeneous morphologies with high variance, obscuring clear statistical significance (**Fig. 2**; **Fig. 6a-c**). Along with the decrease in size, immunostaining revealed that 22q11DS rosettes also exhibited decreased expression of pHH3 and N-cadherin, markers of neuroepithelial identity observed in control rosettes (**Fig. 6e,f**). Specifically, 22q11DS rosettes had, on average, half the number of pHH3+ cells within each rosette (**Fig. 6g**), suggesting that the smaller rosette size may be correlated with decreased neuroepithelial proliferation. In terms of self-organization, most of the 22q11DS rosettes formed in HELP contained a single focal point at the apical center where N-cadherin and actin are localized rather than the open lumen structure that is characteristic of control neural rosettes (**Fig. 6e,f,h**). This morphological difference is significant as lumen formation has been demonstrated to be critical for floor plate patterning in human neural tube organoids.^31^ Collectively, these data demonstrate that within HELP gels, 22q11DS neural rosettes exhibit morphological differences indicative of aberrant proliferation and organization, whereas such differences were not discernable in Matrigel.

**Figure 6.**
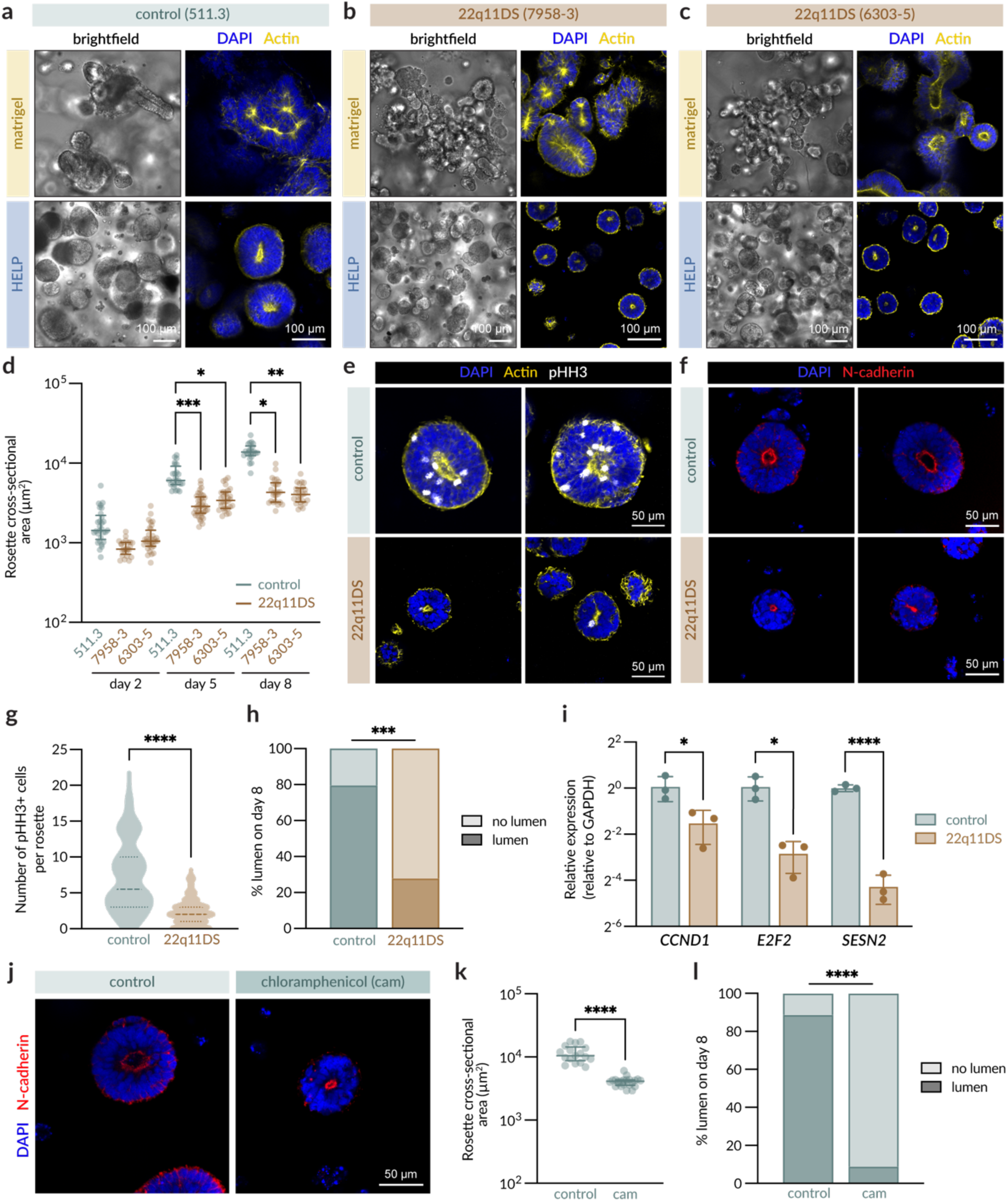
3D Single-lumen neural rosettes derived from 22q11DS patients exhibit altered proliferation and lumen morphologies. **a-c.** Representative brightfield (left) and fluorescence images (right) at day 8 of control (a), 22q11DS patient 1 (b), and 22q11DS patient 2 (c) neural rosettes in Matrigel (top) and Dyn. Slow HELP (bottom) showing nuclei (DAPI, blue) and F-actin (phalloidin, yellow). **d.** Quantification of rosette cross-sectional area at days 2, 5, and 8 in Dyn. Slow HELP (*N* = 3 independent experimental replicate hydrogels; 511.3, *n* = 22-29 rosettes; 7958-3, *n* = 24-37 rosettes; 6303-5, *n* = 25-32 rosettes; data are medians with interquartile range). **e.** Images at day 8 in Dyn. Slow HELP labeled with pHH3 (white) and counterstained for nuclei (DAPI, blue) and F-actin (phalloidin, yellow). **f.** Images at day 8 in Dyn. Slow HELP labeled with N-cadherin (red) and counterstained for nuclei (DAPI, blue). **g.** Quantification of the number of pHH3+ cells per rosette at day 8 (*N* = 3 independent experimental replicate hydrogels; control, *n* = 64 rosettes; 22q11DS, *n* = 64 rosettes; data are medians with interquartile range). **h.** Percentage of rosettes with an open lumen vs. a single central point (*N* = 3 independent experimental replicate hydrogels; data are means). **i.** mRNA expression of *CCND1*, *E2F2*, and *SESN2* in day 8 neural rosettes (*N* = 3 independent experimental replicate hydrogels; data are means ± standard deviation). **j.** Images at day 8 of control neural rosettes in Dyn. Slow HELP treated with 100 µg/ml chloramphenicol (cam) from days 1 to 3 labeled with N-cadherin (red) and counterstained for nuclei (DAPI, blue). **k.** Quantification of rosette cross-sectional area at day 8 after cam treatment (*N* = 3 independent experimental replicate hydrogels; 511.3, *n* = 22-29 rosettes; control, *n* = 17 rosettes; cam, *n* = 22 rosettes; data are medians with interquartile range). **l.** Percentage of rosettes with an open lumen vs. a single central point after cam treatment (*N* = 3 independent experimental replicate hydrogels; data are means). Statistical analyses were Kruskal-Wallis with Dunn’s multiple comparisons test (d), Mann-Whitney test (g, l), and unpaired t-test (h, i, k). *p < 0.05, **p < 0.01, ***p < 0.001, and ****p < 0.0001.

We next validated whether 3D single-lumen neural rosettes derived from patient-derived 22q11DS iPSCs exhibited pathobiological features reported in non-human 22q11DS models. The decreased neural progenitor cell expansion observed in our human-specific cultures is consistent with mouse and zebrafish models of 22q11DS.^66,71^ Within the 22q11.2 deleted region, several genes have been linked to altered neural progenitor cell proliferation and expansion. These include *DGCR8*,^72,73^ *RANBP1*,^66,69^ *CDC45*,^66^ and *PRODH*,^71^ which we confirmed are downregulated in the 22q11DS cultures used here (**Supplementary Fig. 8c**). In particular, *RANBP1* and *CDC45* are highly expressed during development in the ventricular and subventricular zones and are thought to regulate the cell cycle.^66^ In a mouse model of 22q11DS, a previous study identified altered expression of cell cycle regulatory genes within the embryonic mouse cortex.^66^ Therefore, we evaluated whether the expression of these cell cycle related genes beyond the 22q11.2 deleted region were also impaired within our system. Indeed we observed reduced expression of *CCND1*, the gene that encodes cyclin D1, a key regulator of the G1-to-S phase transition of the cell cycle;^74^ *E2F2*, the gene that encodes E2F transcription factor 2, which is critical in cell cycle control;^75^ and *SESN2*, which encodes sestrin 2, a stress-responsive protein that regulates cell proliferation^76^ (**Fig. 6i**). In combination with the decrease in pHH3+ cells, these data suggest that cell cycle regulation is impaired in human 22q11DS cultures, leading to reduced neuroepithelial expansion.

Defective mitochondrial function has also previously been reported to impact neurodevelopment and alter brain size and neural progenitor proliferation *in vivo*, including in 22q11DS animal models.^77,78^ *MRPL40* and *PRODHA* are two 22q11DS genes that encode mitochondrial proteins, and zebrafish single-gene mutants for either gene displayed reduced brain volume, or microcephaly.^71^ Furthermore, *in vivo* treatment with an inhibitor of mitochondrial protein synthesis, chloramphenicol (cam), was found to phenocopy the zebrafish mutants.^71^ Therefore, we examined whether disrupting mitochondrial function *in vitro* within our human, single-lumen neural rosette model would contribute to the morphological phenotypes observed in our 22q11DS rosettes. Control iPSC cultures were treated with cam during the first two days of neural rosette formation. By day 8, the cam-treated neural rosettes were much smaller in size and had similar morphology to the 22q11DS rosettes (**Fig. 6j,k; Supplementary Fig. 9a**), with the majority containing a single central point rather than an open lumen (**Fig. 6l**). Importantly, cam treatment did not impact formation efficiency or rosette circularity (**Supplementary Fig. 9b,c**). These results provide human data to support the idea that mitochondrial function is important for regulating neuroepithelial expansion and early tissue morphogenesis.

Taken together, our human early developmental model of 22q11DS replicates many of the key features of previously reported animal models, including a decrease in pHH3+ proliferative cells, altered expression of cell cycle regulatory genes, and recapitulation of the 22q11DS phenotype upon pharmacological inhibition of mitochondrial function. This suggests that our single-lumen neural rosettes provide a physiologically relevant platform for generating patient-specific avatars to detect disease-related phenotypes and hold potential for future mechanistic and therapeutic studies of early neurodevelopmental disease.

### Outlook

Here, we establish an engineered hydrogel platform for the robust and reproducible generation of single-lumen neuroepithelial tissues. By exploiting cell–matrix interactions to recreate dynamic tissue boundaries, we demonstrate how hydrogel-imposed boundary conditions can guide living tissue assemblies to achieve stable morphogenetic outcomes without artificial top-down constraints. Compared to Matrigel, we demonstrate that defined boundary conditions in a synthetic hydrogel greatly improve reproducibility in tissue morphogenesis, which is useful for probing cell–matrix mechanobiology as well as unveiling disease phenotypes at scale. To date, most previous approaches to generate neural organoids involve aggregating multiple cells together or selecting multicellular structures as a starting point for neural induction.^20,25,27,42,45^ With this approach, the unique ability to form 3D single-lumen neural rosettes from a single iPSC opens up new experimental approaches to probe how individual genetic perturbations affect large-scale organoid morphogenesis.

This work demonstrates how dynamic hydrogel boundary conditions can guide self-organization of early neural tissue structures *in vitro*, establishing design principles that may be applicable to other self-emerging systems. We find that the emergence and organization of 3D single-lumen neural rosettes is highly regulated by matrix biophysical cues. The efficacy of the matrix requires integrin-binding epitopes that promote cell adhesion and signaling, which underlie the establishment of neuroepithelial polarity. Importantly, our results also demonstrate the need for a balance between matrix stress relaxation and matrix-imposed confinement. The material must relax sufficiently to permit multicellular volumetric expansion but maintain enough mechanical resistance to balance the tension exerted by cells on the matrix. Thus, tensional homeostasis, mediated by integrin-induced mechanosignaling, is necessary for the early steps of single-lumen neural rosette formation, including cell survival, lumenogenesis, and polarity acquisition. While the idea of tensional homeostasis originated from studies in cancer biology,^79^ this concept has not been as explored in a developmental context. Our results complement previous work investigating epithelial polarity with MDCK cells, in which tension-based homeostasis was altered through tuning effective ligand concentration in supramolecular hydrogels.^80^ Thus, our *in vitro* hydrogel-enabled platform for single-lumen neuroepithelium formation has the potential to shed light on key developmental requirements *in vivo*. Additional analyses of the dynamic forces exerted during the process of neural tube formation *in vivo* would provide greater insights into the biophysics of neuroepithelial lumen expansion.

Lastly, we demonstrate that 3D single-lumen neural rosettes formed within matrix-defined boundary conditions can robustly manifest disease-related differences in early neurulation, enabling future mechanistic studies to explore the roles of specific genes in regulating human neurodevelopmental phenotypes. Furthermore, we anticipate that this robust and scalable system would facilitate high-throughput screening of drugs for neurodevelopmental disorder-associated phenotypes. In this current work, we have limited the culture period to the initial 8 days to focus on the earliest stages of polarity acquisition and single lumen morphogenesis. Further investigation is still required to explore later stages of neurodevelopment, and future studies will focus on subsequent regionalization and maturation of organoids following neural induction. Taken together, we present a new, matrix-inspired approach to generating 3D single-lumen neural rosettes that has widespread applicability in developmental biology, disease modeling, and genetic perturbation studies.

## Materials and Methods

### ELP expression

ELP was recombinantly expressed in BLR(DE3) *Escherichia coli* (Sigma Aldrich). Briefly, the cells were transformed with a pET15b plasmid containing the appropriate ELP sequence and an ampicillin resistance gene. A single transformed colony was selected and grown overnight at 37 °C, shaking, in Terrific Broth (Thermo Fisher) supplemented with 0.4% glycerol (Thermo Fisher) and 200 µg/ml ampicillin (Thermo Fisher). After 16 h, 20 ml of the starter culture was transferred into each of twelve baffled flasks containing the same growth medium. The cultures were incubated at 30 °C for 24 h, shaking, after which they were collected by centrifugation. The resulting pellet was solubilized in TEN buffer: 10 mM Tris Base (Thermo Fisher), 100 mM sodium chloride (NaCl; Thermo Fisher), 1 mM EDTA (Thermo Fisher); pH 8.0. The resuspended lysate then underwent three freeze–thaw cycles, adding 10 µM deoxyribonuclease I (Sigma Aldrich) and 1 mM phenylmethanesulfonyl fluoride (PMSF; Thermo Fisher) after the first thaw.

ELP was purified through three rounds of thermal cycling, which involves a series of cold spins followed by hot spins. For each round, the pH of the solution was first adjusted to 9.0. The sample was then centrifuged at 15,000 x g for 1 h at 4 °C. For the RGD and R<u>DG</u> variants, 1 M NaCl was added to the supernatant and incubated at 37 °C for 3 h, shaking. For the YIGSR variant, 0.5 M ammonium sulfate (Sigma Aldrich) was used instead of NaCl. The sample was then centrifuged at 15,000 x g for 1 h at 37 °C. The resulting pellet was allowed to re-solubilize in chilled ultrapure water overnight at 4 °C. After the three rounds, one final cold spin was performed, and the resulting supernatant was then dialyzed against chilled ultrapure water for three days, sterile filtered to remove precipitates, and lyophilized.

### Synthesis of hydrazine-functionalized ELP

ELP was modified with hydrazine functional groups by an amidation reaction as previously described.^39^ Briefly, lyophilized ELP was fully dissolved in anhydrous dimethyl sulfoxide (DMSO; Sigma Aldrich) before adding an equal volume of anhydrous *N,N*-dimethylformamide (DMF; Sigma Aldrich) to reach a final concentration of 3.6 wt%. Separately, tri-Boc-hydrazinoacetic acid (2.1 eq. per ELP amine; Sigma Aldrich) was dissolved in the same volume of DMF that was used to dissolve the ELP. Hexafluorophosphate azabenzotriazole tetramethyl uronium (HATU; 2 eq. per ELP amine; Sigma Aldrich) and 4-methylmorpholine (5 eq. per ELP amine; Sigma Aldrich) were then added to activate the tri-Boc-hydrazinoacetic acid for 10 min, stirring. Once activated, the tri-Boc-hydrazinoacetic acid solution was then transferred dropwise to the dissolved ELP. The reaction was allowed to proceed for 20-24 h, stirring. The following day, the ELP was precipitated in ice-cold diethyl ether (Thermo Fisher), centrifuged, and dried under nitrogen gas. To remove the Boc groups, the dried ELP pellet was dissolved in a 1:1 mixture of dichloromethane (DCM; Sigma Aldrich) and trifluoroacetic acid (TFA; Sigma Aldrich). Once dissolved, 5% (v/v) triisopropylsilane (Sigma Aldrich) was added, and the reaction was allowed to proceed for 4 h, stirring. The ELP was then precipitated in ice-cold diethyl ether, centrifuged, and dried under nitrogen gas. This pellet was then dissolved in chilled ultrapure water overnight at 4 °C, dialyzed against chilled ultrapure water for three days, sterile filtered, and lyophilized.

### Synthesis of norbornene-functionalized ELP

ELP was first modified with a PEG-amine by an amidation reaction as previously described.^33^ Briefly, lyophilized ELP was fully dissolved in anhydrous DMSO before adding an equal volume of anhydrous DMF to reach a final concentration of 3.6 wt%. Separately, t-Boc-*N*-amido-PEG12-acid (2.1 eq. per ELP amine; BroadPharm) was dissolved in the same volume of DMF that was used to dissolve the ELP. HATU (2 eq. per ELP amine) and 4-methylmorpholine (5 eq. per ELP amine) were then added to activate the t-Boc-*N*-amido-PEG12-acid for 10 min, stirring. Once activated, the t-Boc-*N*-amido-PEG12-acid solution was then transferred slowly to the dissolved ELP over the course of 10 min. The reaction was allowed to proceed for 20-24 h, stirring. The following day, the ELP was precipitated in ice-cold diethyl ether, centrifuged, dried under nitrogen gas, and deprotected as above. The resulting pellet was then dissolved in chilled ultrapure water overnight at 4 °C, dialyzed against chilled ultrapure water for three days, and lyophilized.

To achieve norbornene-functionalized ELP, the lyophilized ELP-PEG12-amine intermediate was fully dissolved in anhydrous DMSO before adding an equal volume of anhydrous DMF to reach a final concentration of 3.6 wt%. Separately, exo-5-norbornene carboxylic acid (3 eq. per ELP amine; Sigma Aldrich) was dissolved in the same volume of DMF that was used to dissolve the ELP. HATU (3.3 eq. per ELP amine) and 4-methylmorpholine (5 eq. per ELP amine) were then added to activate the exo-5-norbornene carboxylic acid for 10 min, stirring. Once activated, the exo-5-norbornene carboxylic acid solution was then transferred slowly to the dissolved ELP over the course of 10 min. The reaction was allowed to proceed for 20-24 h, stirring. The following day, the ELP was precipitated in ice-cold diethyl ether, centrifuged, and dried under nitrogen gas. The resulting pellet was then dissolved in chilled ultrapure water overnight at 4 °C, dialyzed against chilled ultrapure water for three days, sterile filtered, and lyophilized.

### Synthesis of aldehyde- and benzaldehyde-functionalized HA

HA was first modified with alkyne groups via carbodiimide coupling chemistry as previously described.^81^ Briefly, 100 kDa sodium hyaluronate (HA; LifeCore) was fully dissolved at 1 wt% in MES buffer (0.2 M MES hydrate (Sigma Aldrich), 0.15 M NaCl; pH 4.5). Propargylamine (0.8 eq. per HA dimer unit; Sigma Aldrich) was added and the pH was adjusted to 6.0. N-hydroxysuccinimide (NHS; 0.8 eq. per HA dimer unit; Thermo Fisher) and 1-ethyl-3-[3-dimethylaminopropyl]carbodiimide hydrochloride (EDC; 0.8 eq. per HA dimer unit; Thermo Fisher) were then sequentially added to the reaction mixture. The reaction was allowed to proceed overnight, stirring. The following day, the HA reaction mixture was dialyzed against ultrapure water for three days, sterile filtered to remove precipitates, and lyophilized.

The HA-alkyne intermediate was further modified to yield either aldehyde- or benzaldehyde-functionalized HA via copper catalyzed azide alkyne cycloaddition. Lyophilized HA-alkyne was fully dissolved at 1 wt% in 10x isotonic phosphate buffered saline (PBS; 81 mM sodium phosphate dibasic (Thermo Fisher), 19 mM sodium phosphate monobasic (Thermo Fisher), 60 mM NaCl; pH 7.4) supplemented with 1 mg/ml β-cyclodextrin (Sigma Aldrich). Once dissolved, the solution was degassed under nitrogen for 30 min. In separate vials, 10x solutions of 2.4 mM copper sulfate pentahydrate (Sigma Aldrich) and 45.2 mM sodium ascorbate (Sigma Aldrich) were each prepared in ultrapure water and degassed under nitrogen for 30 min. The sodium ascorbate solution and copper sulfate solution were then added sequentially to the dissolved HA using a syringe, to reach final concentrations of 4.52 mM and 0.24 mM, respectively. For aldehyde functionalization, Ald-CH2-PEG3-Azide (2 eq. per alkyne; BroadPharm) was dissolved in anhydrous DMSO and added to the reaction mixture. For benzaldehyde functionalization, a small-molecule azidobenzaldehyde (2 eq. per alkyne; synthesis previously described^49^) was used instead. The reaction mixture was then degassed under nitrogen for 10 more min, parafilmed, and covered with aluminum foil. The reaction was allowed to proceed overnight, stirring. The following day, an equal volume of 50 mM EDTA (pH 7.0) was added to the reaction and allowed to stir for 1 h. The solution was then dialyzed against ultrapure water for three days, sterile filtered, and lyophilized.

### Synthesis of tetrazine-functionalized HA

HA was modified with tetrazine functional groups via carbodiimide coupling chemistry as previously described.^33^ Briefly, 100 kDa HA was fully dissolved at 1 wt% in 0.1 M MES buffer (pH 7.0). 1-hydrozybenzotriazole hydrate (HOBt; 0.55 eq. per HA dimer unit; Sigma Aldrich) was added and allowed to dissolve for 15 min. Separately, tetrazine amine (0.275 eq. per HA dimer unit; Conju-Probe) was dissolved in a 5:1 mixture of acetonitrile (MeCN; Sigma Aldrich) and ultrapure water. Once dissolved, EDC (0.55 eq. per HA dimer unit) was added and allowed to dissolve. The tetrazine amine solution was then transferred slowly to the dissolved HA over the course of 30 min. The reaction was allowed to proceed overnight, stirring. The solution was then dialyzed against 10% MeCN for two days, followed by ultrapure water for three days, sterile filtered, and lyophilized.

### HELP gel formation

The day prior to HELP gel formation, lyophilized ELP and lyophilized HA were separately dissolved in 10x isotonic PBS overnight at 4 °C to 2 wt% each. Custom-fabricated polydimethylsiloxane (PDMS) molds were prepared by using a biopsy punch to punch out 4-mm diameter circles from a 0.8-mm thick PDMS sheet (Electron Microscopy Sciences). Individual circles were cut out, plasma bonded onto No. 2 glass coverslips, and autoclaved. On the day of gel formation, individual molds were placed into each well of a 24-well culture plate and rinsed with PBS. Dissolved solutions of ELP and HA were kept on ice for the duration of the process, as ELP is sensitive to temperature changes. For Dynamic HELP gels, 5 µl of the dissolved HA solution were pipetted into each mold, and the plate was then placed on ice. Subsequently, 5 µl of the dissolved ELP solution were added to each mold, and the ELP and HA were mixed by swirling the pipette in circles. The plate was then flipped upside down and incubated at room temperature (RT) for 15 min, followed by 37 °C for 10 min. For Static HELP gels, equal volumes of the dissolved HA solution and the dissolved ELP solution were mixed in an Eppendorf tube on ice. In quick succession, 10 µl of the hydrogel mixture were dispensed into each prepared mold. The plate was then incubated on ice for 5 min, then flipped upside down and incubated at RT for 15 min, followed by 37 °C for 10 min.

### Rheological measurement

All rheological measurements were performed with a 20-mm cone-and-plate geometry on a stress-controlled ARG2 rheometer (TA Instruments). Hydrogels of 48 µl were cast directly on the rheometer stage, and a layer of mineral oil was used to maintain sample hydration. Time sweeps were performed during sample gelation at 1% oscillatory strain and 1 rad/s angular frequency. After the plateau modulus was reached, frequency sweeps were performed from 0.1 to 100 rad/s at 1% oscillatory strain and 37 °C. This was followed by a 5 min time sweep at 1% oscillatory strain and 1 rad/s angular frequency for sample equilibration. For stress relaxation measurements, a constant strain of 10% was applied at 37 °C and the resulting stress was measured for a period of 12 h.

### Human iPSC culture, encapsulation, and generation of single-lumen neural rosettes

Human induced pluripotent stem cells (iPSCs) from four healthy patient donors (511, SUN004, 1208, 2242) and two 22q11DS patient donors (6303, 7958) were maintained on Matrigel-coated plates (Corning) in mTeSR Plus medium (Stemcell Technologies). For maintenance passaging, cells were dissociated using Versene (Thermo Fisher). To prepare iPSCs for encapsulation, cells were dissociated into a single cell suspension using Accutase (Thermo Fisher), counted, and resuspended in the dissolved ELP solution at 2x the final desired cell seeding concentration (*i.e.*, for a final concentration of 5,000 cells/µl, cells would be resuspended in ELP at 10,000 cells/µl). Individual hydrogels were formed as described in *HELP gel formation*. After the incubation period, 650 µl of mTeSR Plus supplemented with 10 µM Y27632 (Selleckchem) were added to each well. The next day, the medium was replaced with fresh mTeSR Plus without Y27632. On day 2, the medium was replaced with N3 medium, which consists of DMEM/F12 (Thermo Fisher), Neurobasal (Thermo Fisher), B27 supplement (1:50; Thermo Fisher), N2 supplement (1:100, Thermo Fisher), GlutaMAX (1:100, Thermo Fisher), MEM NEAA (1:100, Thermo Fisher), and Penicillin-Streptomycin (1:100, Thermo Fisher), supplemented with 10 µM SB-431542 (Tocris) and 100 nM LDN-193189 (Stemgent). Medium changes were performed daily.

### Neural induction of iPSCs in suspension culture

Human iPSCs were dissociated into a single cell suspension using Accutase and counted. 1.5 million cells were added to one well of an AggreWell^TM^800 24-well plate (Stemcell Technologies) in mTeSR Plus medium supplemented with Y27632 and centrifuged at 100 x g for 3 min to create 5,000 cell aggregates. The next day, a media change was performed very carefully with fresh mTeSR Plus without Y27632. On day 2, the spheroids were transferred from the microwells using a cut p1000 pipette tip onto a 70-µm cell strainer such that any single cells or debris would flow through. The strainer was inverted over a 50-ml conical tube and 12 ml of E6 medium (Thermo Fisher) supplemented with 10 µM SB-431542 and 100 nM LDN-193189 was flowed through to collect the spheroids. The entire solution was then transferred to an ultra-low attachment culture dish. Medium changes were performed daily.

### Neural induction of iPSCs in Matrigel

Growth factor-reduced Matrigel (Corning) was thawed on ice. Human iPSCs were dissociated into a single cell suspension using Accutase and counted. To prepare iPSCs for encapsulation, cells were dissociated into a single cell suspension using Accutase, counted, and resuspended on ice in the thawed Matrigel solution at 1,000 cells/µl. Then, 10 µl of the cell suspension were dispensed directly into each prepared mold. The plate was flipped upside down and incubated at RT for 15 min, followed by 37 °C for 10 min, flipping the plate every 5 min. After the incubation period, 650 µl of mTeSR Plus supplemented with 10 µM Y27632 were added to each well. The next day, the medium was replaced with fresh mTeSR Plus without Y27632. On day 2, the medium was replaced with N3 medium supplemented with 10 µM SB-431542 and 100 nM LDN-193189. Medium changes were performed daily.

### Viability and proliferation assays

Cell viability was evaluated by incubating hydrogels in PBS containing 2 µM calcein-AM (Invitrogen) and 4 µM ethidium homodimer-1 (Invitrogen) for 10 min at 37 °C. Following incubation, samples were rinsed with PBS and imaged using a confocal microscope (Leica SPE). Metabolic activity was measured with the alamarBlue Cell Viability Reagent (Invitrogen) according to the manufacturer’s protocol. Cell proliferation was quantified by measuring double-stranded DNA content using the Quant-iT PicoGreen dsDNA Assay Kit (Invitrogen) according to the manufacturer’s protocol. To obtain cell lysates, the hydrogels were first digested in an equal volume of PBS supplemented with hyaluronidase (2000 U/ml; Sigma Aldrich) and elastase (250 U/ml; Sigma Aldrich) for 20 min at 37 °C. Then, 500 µl of cell lysis buffer (20 mM Tris-HCl (Sigma Aldrich), 150 mM NaCl, 0.5% Triton X-100 (Sigma Aldrich); pH 7.4) was added, and the samples were incubated on ice for 20 min. Samples were stored at −80 °C until further analysis.

### Immunostaining

For HELP gels, samples were fixed in 4% paraformaldehyde (PFA; Electron Microscopy Sciences) in PBS for 20 min at RT. For spheroids cultured in suspension, individual spheroids were transferred to separate wells of a 96-well plate and fixed in 4% PFA in PBS overnight at 4 °C. For Matrigel cultures, samples were fixed in 4% PFA and 0.1% glutaraldehyde (Sigma Aldrich) in PBS for 20 min at RT. After fixation, samples were washed three times with PBS for 10 min each. Samples were permeabilized with PBS-T (0.25% Triton X-100 in PBS) for 1 h and blocked with 0.5% Triton X-100, 5% goat serum (Thermo Fisher), and 5% (w/v) bovine serum albumin (BSA; Roche) in PBS for 3 h at RT. Primary antibodies were diluted in antibody dilution solution, which consists of 0.5% Triton X-100, 2.5% goat serum, and 2.5% (w/v) BSA in PBS. Primary antibodies used include: rabbit anti-Sox2 (1:200, Cell Signaling, 23064), mouse anti-N-cadherin (1:200, BD Biosciences, 610920), rabbit anti-Pax6 (1:200, Cell Signaling, 60433), mouse anti-ZO-1 (1:200, Invitrogen, 33-9100), mouse anti-Nestin (1:200, Millipore, MAB326), mouse anti-pHH3 (1:200, Santa Cruz, sc-374669), mouse anti-βIII-tubulin (1:200, Cell Signaling, 4466), mouse anti-PKCζ (1:50, Santa Cruz, sc-17781), mouse anti-FAK (1:200, Protein Tech, 66258-1-Ig). Samples were incubated in the primary antibody solution for 48 h at 4 °C, rocking. After incubation, samples were washed three times with PBS-T for 30 min each and then incubated in antibody dilution solution with 4′,6-diamidino-2-phenylindole (DAPI; 1 μg/mL, Cell Signaling, 4083s), TRITC-phalloidin (0.2 µg/ml, Sigma Aldrich, P1951), and the appropriate secondary antibodies overnight at 4 °C, rocking. Secondary antibodies used include: goat anti-rabbit Alexa Fluor 488 (1:500, Invitrogen, A-11008) and goat anti-mouse Alexa Fluor 647 (1:500, Invitrogen, A-21236). After incubation, samples were washed three times with PBS for 30 min each and imaged.

### Microscopy

Live brightfield imaging of iPSCs in HELP was performed on a Leica THUNDER Imager with a 10x air objective. Brightfield images of neural rosettes and fluorescence images of mScarlet- and GFP-expressing iPSCs were taken on a Leica THUNDER Imager with either 10x or 20x air objectives. Fluorescence images of immunostained samples were taken on a Leica SP8, Leica SPE, or Zeiss LSM 780 confocal microscope with either 20x air, 40x oil, or 63x oil objectives. Spheroids were placed in EasyIndex optical clearing solution (LifeCanvas Technologies) on a glass coverslip for imaging.

### RNA isolation and qPCR

To release the encapsulated cells, hydrogels were first digested in an equal volume of PBS supplemented with hyaluronidase (2000 U/ml; Sigma Aldrich) and elastase (250 U/ml; Sigma Aldrich) for 20 min at 37 °C. Then, 500 µl of TRIzol reagent (Invitrogen) were added for each hydrogel, and the samples were immediately transferred to −80 °C for storage. When ready for RNA isolation, samples were thawed on ice and mechanically disrupted using a probe sonicator (Heilscher UP50H; 50% amplitude, 25 W, 30 kHz, 0.5 s pulse cycle). RNA was isolated via phenol-chloroform extraction using Phasemaker^TM^ tubes (Invitrogen) and then precipitated with isopropanol (Thermo Fisher). The RNA pellets were then washed twice with 70% ethanol (Thermo Fisher), air-dried, and resuspended in nuclease-free water. Reverse transcription was performed on 500 ng of RNA using the High Capacity cDNA Reverse Transcription Kit (Thermo Fisher). The resulting cDNA was diluted ten-fold with nuclease-free water. Quantitative PCR reactions were prepared by combining 6.6 µl of diluted cDNA with 0.9 µl of a 5 µM forward/reverse primer mixture (**Supplementary Table 1**; Integrated DNA Technologies) and 7.5 µl of Fast SYBR Green Master Mix (Thermo Fisher). The PCR reaction was carried out either on a QuantStudio^TM^ 3 Real-Time PCR System (Applied Biosystems) or QuantStudio^TM^ 6 Flex Real-Time PCR System (Applied Biosystems). Relative gene expression was quantified using the ΔCT method.

### Inhibitor treatment

For studies involving inhibition of mechanosignaling pathways, the drug was added starting on day 1 of culture. The inhibitors used were Y27632 (10 µM; Selleckchem, ROCK inhibitor), blebbistatin (10 µM; Abcam, myosin II inhibitor), ML-7 (25 µM; Tocris, myosin light chain kinase inhibitor), ML-141 (2 µM; Tocris, Cdc42 Rho family inhibitor), CK666 (100 µM; Sigma Aldrich, Arp2/3 inhibitor), and latrunculin A (1 µM; Tocris, actin polymerization inhibitor). For inhibition of mitochondrial protein synthesis, chloramphenicol (Sigma Aldrich) was used at 100 µg/ml and added to the culture media from days 1 to 3.

### Image analysis

Quantification of spheroid or rosette cross-sectional area and circularity were made based on binary masks constructed in Fiji. Masks were obtained by taking the maximum z-projection of the DAPI channel, applying a Gaussian blur, thresholding, then filling the holes. Analyze Particles identified individual spheroids or rosettes, and area and circularity metrics were extracted from the Measure command. For quantification of lumen metrics, individual lumens were manually traced in Fiji based on actin staining, and area and circularity metrics were extracted from the Measure command. For localization of Sox2 and N-cadherin markers, a line was manually drawn across the center of the rosette structure in Fiji. The intensity profile was then plotted along the line for the relevant channels. The intensity and position were normalized such that the maximum intensity was 1, and the edges of the luminal surface were positioned at −1 and +1. The number of rosettes or spheroids formed was quantified using CellProfiler based on a maximum z-projection of the DAPI channel. The image was thresholded and objects with 2D projected areas less than 400 µm^2^ were removed before counting. The percentage of polarized cysts were obtained by manual counting of a z-stack based on DAPI and actin staining. For manual counting, sample identities were blinded to minimize bias.

### Statistical analysis and reproducibility

The data collected for this manuscript were obtained from over three expressions and modifications of ELP and three modifications of HA. Results were consistent across all material batches. Statistical analyses for this study were performed using GraphPad Prism version 10 software. Details of specific statistical methods for each figure are included within the figure captions. For all studies, not significant (ns; p > 0.05), *p < 0.05, **p < 0.01, ***p < 0.001, and ****p < 0.0001.

## Acknowledgements

The authors thank K. Loh (Stanford Developmental Biology) and S. Paşca (Stanford Psychiatry and Behavioral Sciences) for providing some human iPSC lines, F. Yang (Stanford Bioengineering) and O. Chaudhuri (Stanford Mechanical Engineering) for use of equipment, and the Stanford Cell Sciences Imaging Facility for microscopy use. M.S.H. acknowledges support from the Sarafan ChEM-H Chemistry/Biology Interface Program as an O’Leary-Thiry Fellow, a National Science Foundation (NSF) Graduate Research Fellowship (DGE-1656518), a National Institutes of Health (NIH) NRSA Pre-doctoral Fellowship (F31-NS132505), and the Stanford University Gerald J. Lieberman Fellowship. J.G.R. acknowledges support from an NSF Graduate Research Fellowship (DGE-1656518) and the Stanford Smith Family Graduate Fellowship. R.S.N. acknowledges support from an NIH K99/R00 MOSAIC Postdoctoral Career Transition Award (K99-HL169844). S.C.H. acknowledges support from the NIH (R01-MH137333), NSF (DMR-2427971), the Stanford Brain Organogenesis Program in the Wu Tsai Neurosciences Institute and Bio-X, and the Stanford Maternal and Child Health Research Institute through the Uytengsu-Hamilton 22q11 Neuropsychiatry Research Award Program. Part of this work was performed at the Stanford Nano Shared Facilities, supported by the NSF (ECCS-2026822).

## Author Contributions Statement

M.S.H. and S.C.H. conceived and initiated the project. M.S.H. designed the research, synthesized the materials, conducted the experiments, and analyzed the data. J.G.R. and T.D.P. provided guidance on experimental design and data interpretation. D.K. assisted with suspension neural organoid culture and characterization. K.P.P. assisted with immunostaining and image analysis. D.P. and T.Y. assisted with experiments pertaining to mechanical inhibition and characterization of 22q11DS cultures. Y.L. assisted with qPCR. R.S.N. assisted with biomaterials synthesis. M.S.H. and S.C.H. wrote the manuscript. S.C.H. supervised the study. All authors edited and approved the manuscript.

