## Supplementary Information for "Hydrogel-imposed boundary conditions guide single-lumen neuroepithelial morphogenesis"

**Affiliations**

**This PDF file includes:**

*Supplementary Figure 1. Mechanical characterization of engineered HELP hydrogel.*

*Supplementary Figure 2. Reproducibility of single-lumen neural rosette formation across multiple cell lines when encapsulated as single cells in HELP hydrogels prior to neural induction.*

*Supplementary Figure 3. Characterization of 3D neural rosettes within HELP hydrogels.*

*Supplementary Figure 4. Rheology of Static, Dynamic Slow, and Dynamic Fast HELP hydrogels.*

*Supplementary Figure 5. Cell density-dependence of neural rosette formation in HELP hydrogels.*

*Supplementary Figure 6. Characterization of 3D neural rosettes within Static, Dyn. Slow, and Dyn. Fast HELP hydrogels.*

*Supplementary Figure 7. Neural rosette formation in the presence of inhibitors of mechanosignaling pathways.*

*Supplementary Figure 8. Additional characterization of 22q11DS single-lumen neural rosette cultures.*

*Supplementary Figure 9. Chloramphenicol treatment does not prevent 3D neural rosette formation.*

*Supplementary Table 1. Primers used for qPCR experiments.*

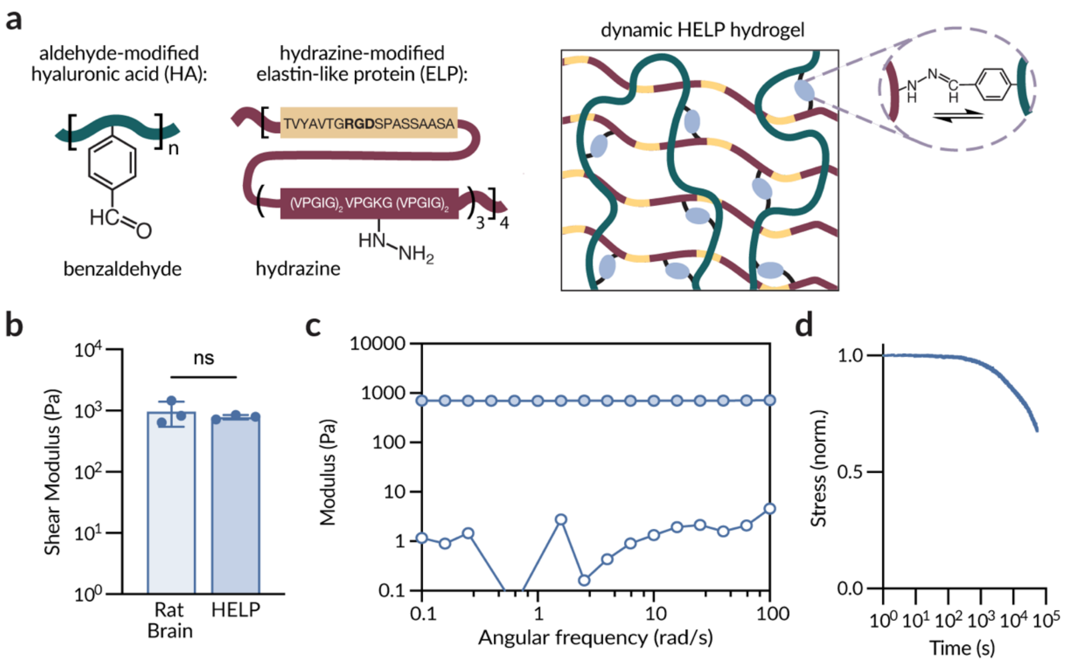


**Supplementary Figure 1.** Mechanical characterization of engineered HELP hydrogel.
**a.** Schematic of HELP hydrogel components: aldehyde-modified hyaluronic acid (HA) and hydrazine-modified elastin-like protein (ELP). The ELP amino acid sequence includes the fibronectin-derived, integrin-adhesive RGD peptide sequence. **b.** Shear storage moduli of rat brain tissue and engineered HELP hydrogel, reported at 1 rad/s. **c.** Frequency sweep of engineered HELP hydrogel at 1% strain showing storage moduli (G', filled symbols) and loss moduli (G", open symbols). **d.** Representative stress relaxation curve of engineered HELP hydrogel under a constant 10% strain.

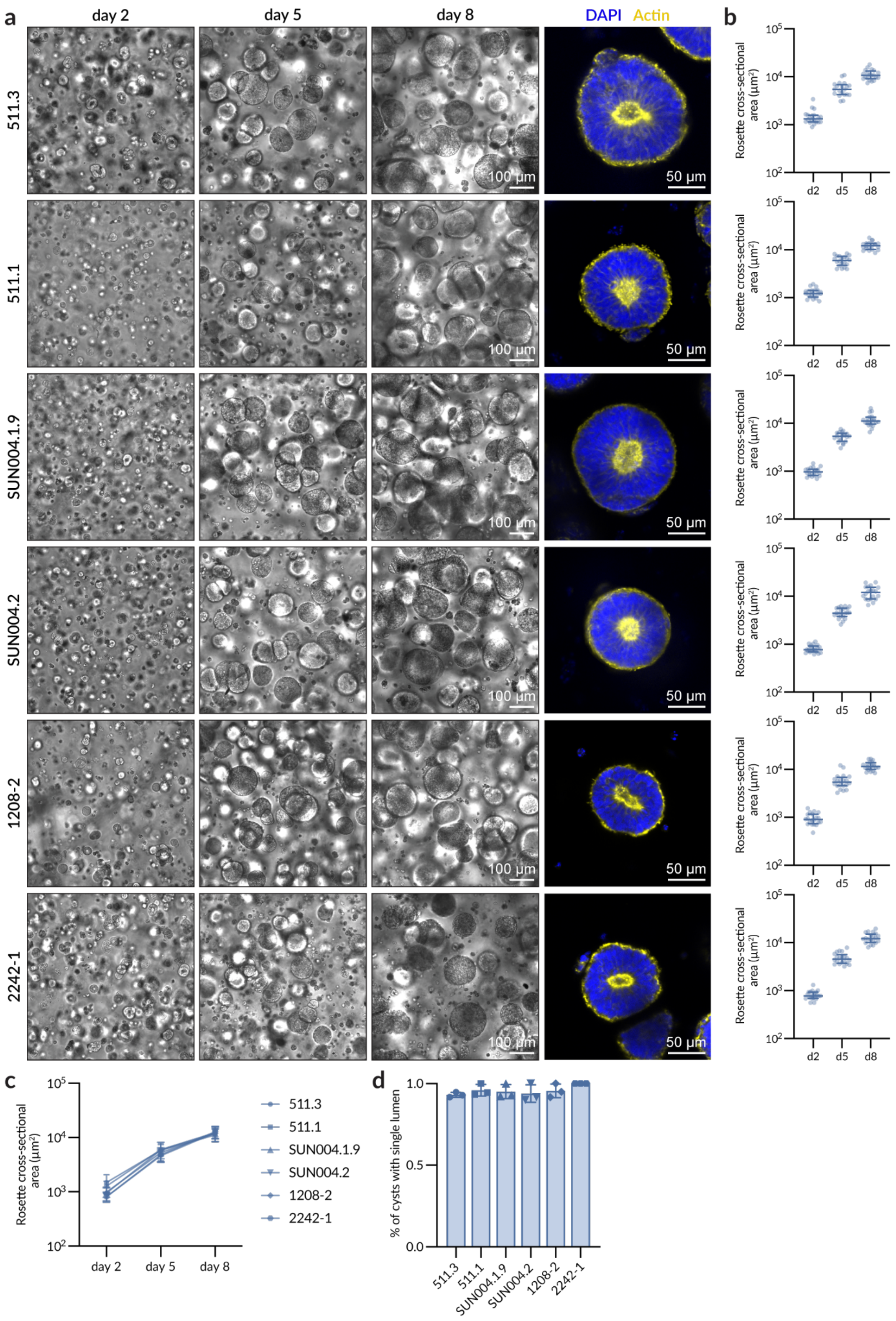


**Supplementary Figure 2.** Reproducibility of single-lumen neural rosette formation across multiple cell lines when encapsulated as single cells in HELP hydrogels prior to neural induction.

**a.** Representative brightfield images at days 2, 5, and 8 (left) and fluorescence images at day 8 (right) of neural rosettes in HELP for multiple iPSC lines. Nuclei are counterstained with DAPI (blue), and F-actin is counterstained with phalloidin (yellow). **b.** Quantification of rosette cross-sectional area at days 2, 5, and 8 (*N* = 3 independent experimental replicate hydrogels; data are medians with interquartile range). **c.** Overlay of rosette growth rates of multiple iPSC lines. **d.** Quantification of the percentage of cysts with a single lumen for multiple iPSC lines (*N* = 3 independent experimental replicate hydrogels; data are means ± standard deviation).

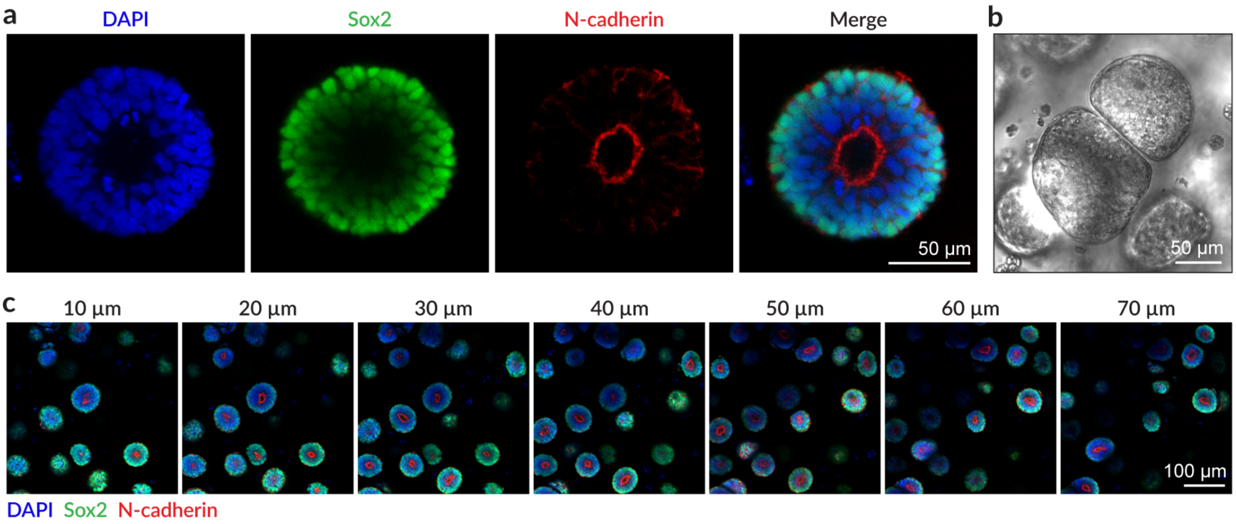


**Supplementary Figure 3.** Characterization of 3D neural rosettes within HELP hydrogels. **a.** Representative fluorescence images at day 8 of neural rosettes in HELP, showing Sox2 expression throughout the rosette, with stronger signal near the basal surface, and a strong ring of N-cadherin at the apical surface. **b.** Representative brightfield image at day 8 of neural rosettes in HELP, showing the lack of fusion between nearby rosettes. **c.** Deconstructed z-stack frames at day 8 of neural rosettes in HELP, showing single-lumen formation is reproducible throughout the entire depth of the hydrogel.

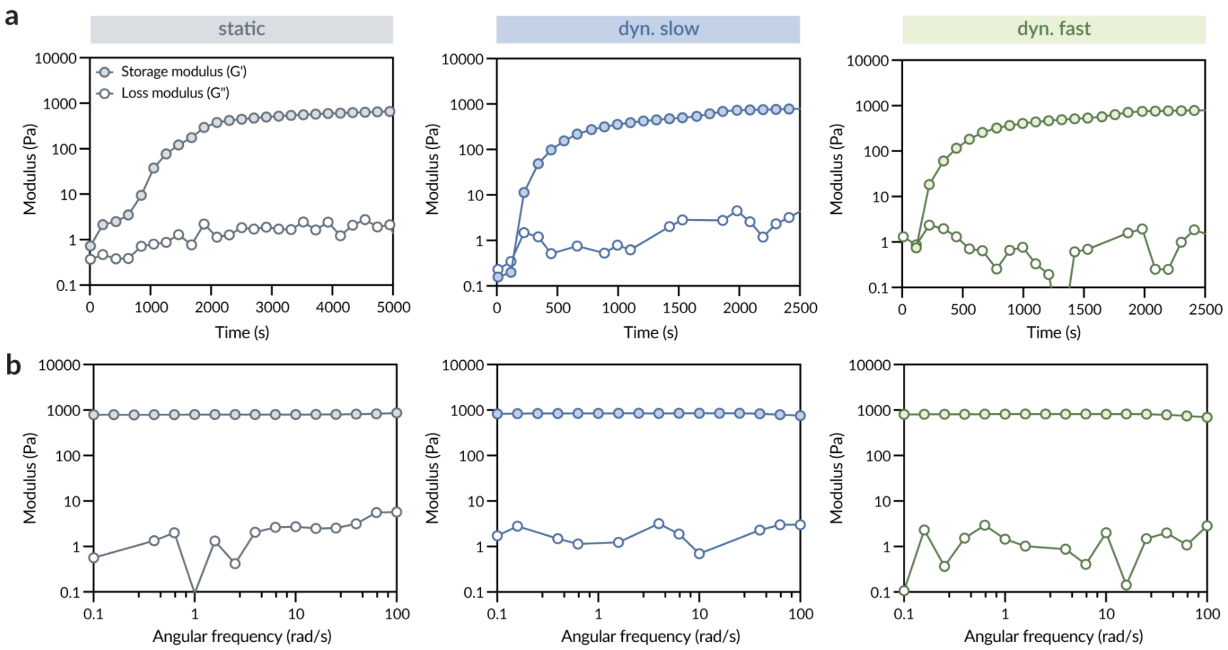


**Supplementary Figure 4.** Rheology of Static, Dynamic Slow, and Dynamic Fast HELP hydrogels.
**a.** Time sweep of HELP hydrogels (Static, Dyn. Slow, and Dyn. Fast) showing storage moduli (G', filled symbols) and loss moduli (G", open symbols) during crosslinking. **b.** Frequency sweep at 1% strain showing storage moduli (G', filled symbols) and loss moduli (G", open symbols) for all three HELP hydrogel formulations.

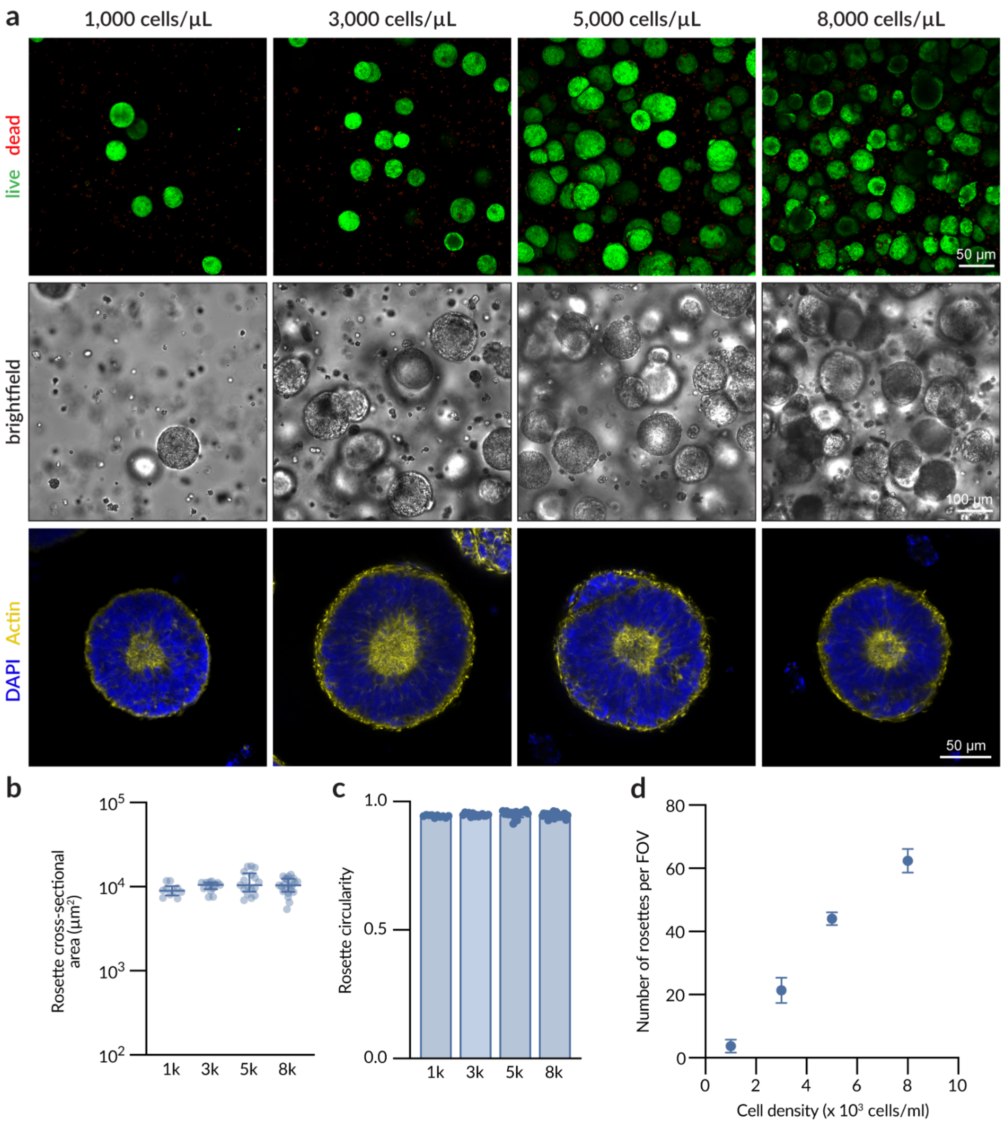


**Supplementary Figure 5.** Cell density-dependence of neural rosette formation in HELP hydrogels.

**a.** Representative fluorescence images labeled with calcein-AM and ethidium homodimer-1 (top), representative brightfield images (middle), and representative fluorescence images labeled with DAPI and actin (bottom) at day 8 of neural rosettes in Dyn. Slow HELP with different initial iPSC seeding densities. **b.** Quantification of rosette cross-sectional area at day 8 (*N* = 3 independent experimental replicate hydrogels; data are medians with interquartile range). **c.** Quantification of rosette circularity at day 8 (*N* = 3 independent experimental replicate hydrogels; data are means ± standard deviation). **d.** Number of rosettes per field of view (FOV; *N* = 3 independent experimental replicate hydrogels; data are means ± standard deviation).

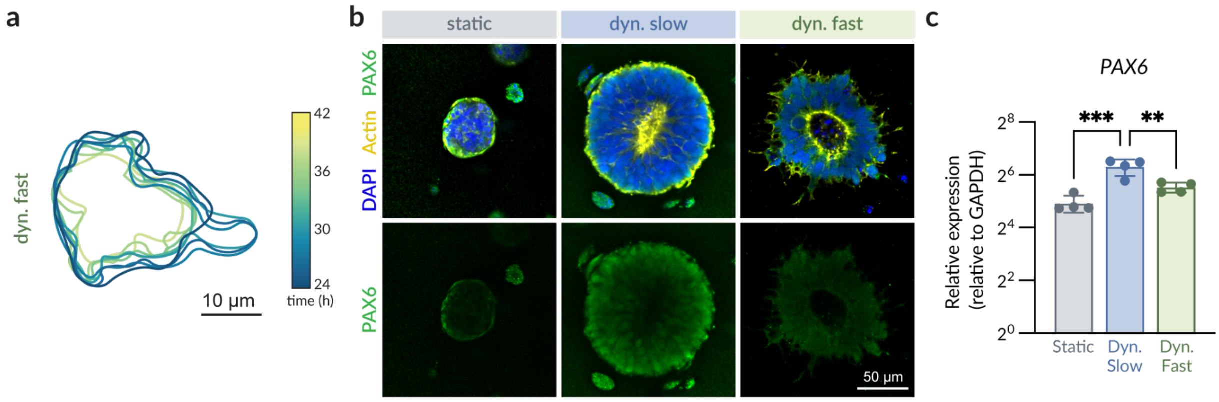


**Supplementary Figure 6.** Characterization of 3D neural rosettes within Static, Dyn. Slow, and Dyn. Fast HELP hydrogels.
**a.** Overlaid outlines of the perimeter of iPSC clusters within Dyn. Fast HELP from 24 h to 42 h of encapsulation showing a spheroid that retracts in size. **b.** Representative fluorescence images at day 8 of neural rosettes in Static, Dyn. Slow, and Dyn. Fast HELP labeled with PAX6 (green) and counterstained for nuclei (DAPI, blue) and F-actin (phalloidin, yellow). **c.** mRNA expression of *PAX6* after 8 days of culture ($N = 4$ independent experimental replicate hydrogels; data are means ± standard deviation). Statistical analyses were one-way ANOVA with Tukey's multiple comparisons test (c). $^{**}p < 0.01$ and $^{***}p < 0.001$.

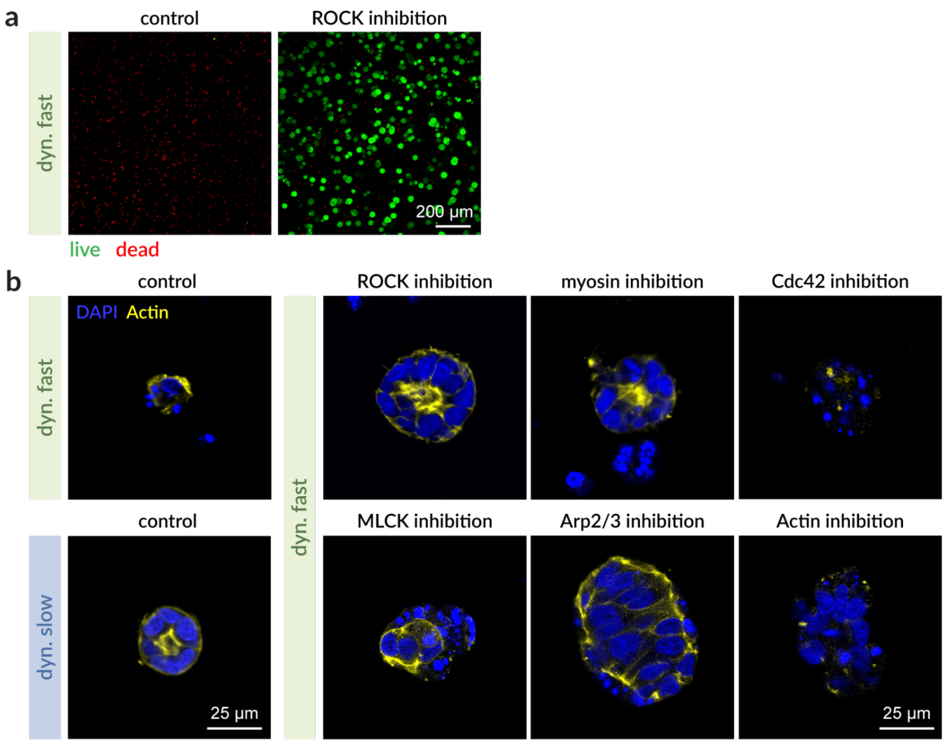


**Supplementary Figure 7.** Neural rosette formation in the presence of inhibitors of mechanosignaling pathways.

**a.** Representative fluorescence images at day 3 of neural rosettes in Dyn. Fast HELP with and without 10 µM Y27632 labeled with calcein-AM (live) and ethidium homodimer-1 (dead). **b.** Representative fluorescence images at day 3 of neural rosettes in Dyn. Fast and Dyn. Slow HELP controls (left panel) and in Dyn. Fast HELP with inhibitors (right panel): 10 µM Y27632, 10 µM blebbistatin, 2 µM ML-141, 25 µM ML-7, 100 µM CK666, and 1 µM latrunculin A showing nuclei (DAPI, blue) and F-actin (phalloidin, yellow).

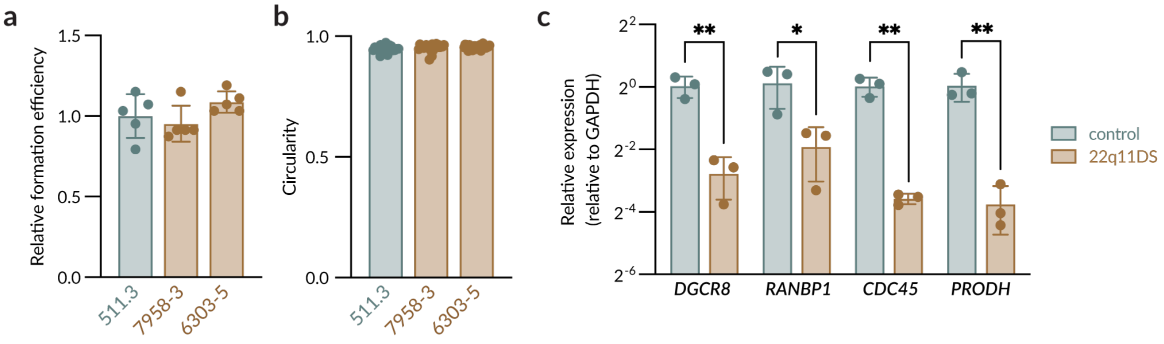


**Supplementary Figure 8.** Additional characterization of 22q11DS single-lumen neural rosette cultures.
**a.** Quantification of 22q11DS spheroid formation efficiency at day 8, normalized to 511.3 control (*N* = 5 independent experimental replicate hydrogels; data are means ± standard deviation). **b.** Quantification of spheroid circularity at day 8 (*N* = 3 independent experimental replicate hydrogels; 511.3, *n* = 22 rosettes; 7958-3, *n* = 25 rosettes; 6303-5, *n* = 25 rosettes; data are means ± standard deviation). **c.** mRNA expression of *DGCR8*, *RANBP1*, *CDC45*, and *PRODH* in day 8 neural rosettes (*N* = 3 independent experimental replicate hydrogels; data are means ± standard deviation). Statistical analyses were one-way ANOVA with Tukey's multiple comparisons test (a, b) and unpaired t-test ©. *p < 0.05 and **p < 0.01.

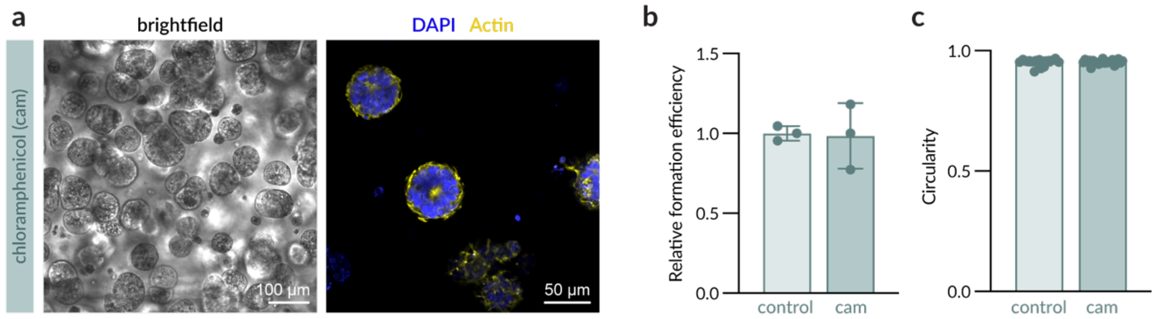


**Supplementary Figure 9.** Chloramphenicol treatment does not prevent 3D neural rosette formation.

**a.** Representative brightfield images (left) and fluorescence images (right) at day 8 of neural rosettes (511.3 line) in Dyn. Slow HELP treated with 100 µg/ml chloramphenicol (cam) from days 1 to 3. Nuclei are counterstained with DAPI (blue), and F-actin is counterstained with phalloidin (yellow). **b.** Quantification of spheroid formation efficiency for cam-treated cultures at day 8, normalized to untreated control (*N* = 3 independent experimental replicate hydrogels; data are means ± standard deviation). **c.** Quantification of spheroid circularity at day 8 (*N* = 3 independent experimental replicate hydrogels; control, *n* = 17 rosettes; cam, *n* = 22 rosettes; data are means ± standard deviation). Statistical analyses were unpaired t-test (b,c).

| Target | Forward Primer (5’ to 3’) | Reverse Primer (5’ to 3’) |
|---|---|---|
| *GAPDH* | AGGTCGGTGTGAACGGATTTG | TGTAGACCATGTAGTTGAGGT |
| *PAX6* | TGCTGCACATTTGGAGACAC | TGCAGAAATGTGCTGCTGTG |
| *MYH9* | CAGCAAGCTGCCGATAAGTAT | CTTGTCGGAAGGCACCCAT |
| *FAK* | GCTTACCTTGACCCCAACTTG | ACGTTCCATACCAGTACCCAG |
| *RAC1* | ATGTCCGTGCAAAGTGGTATC | CTCGGATCGCTTCGTCAAACA |
| *VIM* | GACGCCATCAACACCGAGTT | CTTTGTCGTTGGTTAGCTGGT |
| *DGCR8* | AGCGTGAGCTTTACCGAGAG | CTACCCCGTCACCAACACTC |
| *RANBP1* | AATACAGACGAGTCCAACCATGA | GAACAGTTTTGCCCGCATTTTA |
| *CDC45* | TTCGTGTCCGATTTCCGCAAA | TGGAACCAGCGTATATTGCAC |
| *PRODH* | CCCTGCTTCGGCACTACAG | GGGCCTGGTATTGCTTGTCC |
| *CCND1* | GCTGCGAAGTGGAAACCATC | CCTCCTTCTGCACACATTTGAA |
| *E2F2* | CGTCCCTGAGTTCCCAACC | GCGAAGTGTCATACCGAGTCTT |
| *SESN2* | AAGGACTACCTGCGGTTCG | CGCCCAGAGGACATCAGTG |

**Supplementary Table 1.** Primers used for qPCR experiments.
